# Charting infant sleep cycle development using actigraphy: Longitudinal evidence for cycle lengthening within the first year of life from 35,000 hours of sleep

**DOI:** 10.1101/2025.07.10.664225

**Authors:** Grégory Hammad, Sarah F. Schoch, Max Engelmann, Zoe Spock, Salome Kurth, Eva C. Winnebeck

**Author notes:** The authors contributed equally to the work and share first authorship (Grégory Hammad, Sarah F. Schoch). The authors contributed equally to the work and share last authorship (Salome Kurth, Eva C. Winnebeck).

## Abstract

Sleep is marked by ultradian cycles, coinciding with the alternation of rapid eye movement (REM) and non-rapid eye movement sleep (NREM). These sleep cycles are a striking feature exhibited from infancy through adulthood, yet their underlying mechanisms and functional relevance remain elusive, calling for large-scale longitudinal studies. Here we leverage Locomotor Inactivity During Sleep (LIDS), previously established in adults as an accessible means to track sleep cycles, to chart ultradian sleep cycle dynamics within the first year of life at scale. Specifically, we analyzed *>* 35, 000 hours of sleep from a longitudinal dataset of 152 infants with 10 days of actigraphy at 3, 6, and 12 months of age. Using advanced signal processing techniques, we demonstrate the existence of rhythmic patterns in LIDS in infants with a cycle length of *∼* 60 minutes. Cycles were not only shorter in infants than parents (*−* 11.8 min, 95% C.I. [*−* 13.8, *−* 9.8]) but also increased by *∼* 10 min from 3 to 12 months of age, mirroring previous results in smaller samples for NREM-REM sleep cycles. This increase was found to be partially mediated by an increase in sleep bout duration with age (+1.0 min/h, [+0.9, +1.2]). Longer cycles were also found in infants still breastfed at 12 months (+2.5 min [+0.4, +4.5]) and their breastfeeding mothers (+6.7 min [+0.5, +12.9]). We also explored variations in non-rhythmic LIDS patterns in relation to endogenous and exogenous factors. Overall, our results provide evidence of a link between rhythmic patterns in limb inactivity and sleep cycles in infancy, underscoring the potential of studying these in large-scale population cohorts for developmental and health outcomes.

## Introduction

Ultradian rhythms, biological rhythms with cycle lengths in the hour range and no obvious environmental synchronizers, are manifest across the animal kingdom [1] and across biochemical, physiological and behavioral processes [2, 3] such as gene expression [4, 5], hormone release [6], body temperature [7], feeding habits [8] and motor activity [9, 10]. Despite the ubiquity of ultradian rhythms, their underlying generative mechanisms are largely unknown [11], and their biological function(s) remains equally elusive [12].

Around 70 years ago, in 1955, Aserinsky and Kleitman observed ultradian rhythms in “bodily movements” in infants during sleep, synchronized with the peculiar ultradian rhythms of ocular motility they had previously noticed [13], confirming an even earlier report of this phenomenon [14, 15]. Two years later, Dement and Kleitman described in detail similar observations in adults: brain activity, eye and body movements vary in an ultradian cyclic manner throughout the night [16]. These discoveries led to the first definition of ultradian sleep cycles based on cyclic changes in sleep physiology. Following the shift to an EEG-oriented definition of sleep and sleep stages, Feinberg and Floyd updated this early definition in 1979 [17], which is still in use today: Cycles are defined as the alternation between non-REM (NREM) sleep periods (including all stages of NREM sleep and intermittent wake) and REM sleep periods (REM sleep and intermittent wake over 5 min total) and are accordingly called NREM-REM cycles. A recent retrospective study of *∼* 6000 polysomnographic recordings reported a median sleep cycle length of 96 min in adults with wide variation in cycle length between and within individuals [18].

Despite these early observations of the cyclic nature of sleep, much about the ultradian dynamics of sleep remains elusive today. For example, mechanisms of sleep cycle generation and the functional relevance of rhythm characteristics are still enigmatic (although see recent advances in rodents [19]). This lack of understanding may be largely driven by the difficulty of monitoring sleep cycles at scale, which is necessary to overcome the variability in cycle dynamics characteristic of all ultradian rhythms. Hence, using simpler recording methods than polysomnography, which can be more easily deployed in large numbers of individuals, longitudinally and in everyday environments, appears to be an intuitive path to gaining more data and thus a better understanding of ultradian sleep cycles.

In 2018, we demonstrated that Locomotor Inactivity During Sleep (LIDS), derived from simple wrist-worn actigraphy in everyday life, is an easy method to detect ultradian rhythms in movement during sleep in school-age children, adolescents and adults, and their dynamics are highly similar to NREM-REM sleep cycles assessed with polysomnography [20]. The detected inactivity rhythms oscillate with a cycle length of *∼* 90 *−* 110 min and show characteristic variability, while the mean inactivity levels were found to decline progressively overnight. This decline as well as the oscillation amplitudes were reduced with increasing age, suggesting not only usefulness for ultradian monitoring but also potentially for depth of sleep and homeostatic sleep pressure. Here, we show that the LIDS methodology can also be applied during a particularly critical period in life that is marked by rapid and substantial changes in sleep: infancy.

During the first year of life, sleep undergoes remarkable development: starting from a polyphasic sleep-wake pattern after birth with multiple short sleep bouts scattered almost evenly across the 24-hour day, sleep bouts gradually consolidate, and become increasingly concentrated during the night [21, 22, 23]. This change in sleep-wake organization is attributed to the maturation of two important sleep regulatory systems: the circadian system, which promotes wakefulness during the biological day and sleep during the biological night, and the homeostatic system, which tracks the wake-dependent accumulation of sleep pressure [24, 25].

Interestingly, in early infancy, sleep states are commonly referred to as quiet (NREM) and active (REM) sleep suggesting differences in motor behavior which help demarcate sleep cycles, which also initially led to their discovery by Denisova and by Aserinsky [14, 15, 13]. Only after the first two to three months of life do the differences in EEG appear that clearly distinguish REM and NREM sleep and the NREM substages. This is intriguing as it might indicate that rhythmic motor activity predates the appearance of specific brain activity patterns that allow the classification of sleep epochs into discrete stages. It might also suggest that while sleep physiology is still maturing, the ultradian cycling mechanism is already functional at an early life stage and that its manifestation might be greater in motor activity than in brain activity. Indeed, monitoring of infants’ breathing and body movements with a pressure sensitive mattress [26] revealed cycles in quiet sleep with a period of 63 min at 2 to 5 postnatal weeks. Evidence for the presence of such cycles as early as the first postnatal days was also put forward by Freudigman and Thoman in 1994 [27]. Therefore, already from the first weeks of life, infant sleep alternates between two states [28], a clear manifestation of ultradian sleep cycles. These sleep cycles are on average 50-60 min long [14, 15, 13, 29, 30, 31] and thus markedly shorter than the 90-110-min cycles observed in adults. Studies based on small or cross-sectional samples suggest that cycles may gradually lengthen over the course of infancy and early childhood to reach adult-length cycles by school-age [32]However, strong longitudinal evidence is lacking given the cumbersome nature of polysomnographic recordings especially in this age group.

Here, we used a longitudinal infant and parent dataset with ankle-actigraphy recordings at 3, 6, and 12 months after birth to explore the potential of using motor patterns during sleep via our LIDS methodology to study infant sleep dynamics, particularly those of ultradian sleep cycles. We address multiple questions: Are there ultradian rhythms detectable in infant inactivity during sleep as early as 3 months of age? Do these rhythms show characteristics typical of NREM-REM cycles at this age, i.e., a cycle length around 50-60 min with high variability and shorter lengths than those in adults and their parents? And finally, how do these ultradian rhythms change over the first year of an infant’s life, both in infants and parents? For the latter question, we explored not only cycle lengthening with age but also associations with multiple other key endogenous and exogenous factors including sex, sleep-wake cycle maturation, breastfeeding and sleep location, which are held to influence sleep quality but have never been systematically assessed in large datasets for links with ultradian sleep cycles.

## Methods

### Participants

The work presented here analyzed data from a longitudinal study of infant sleep and gut microbiota [33, 34, 35], in which a total of 162 infants living in the area of Zurich were enrolled. In addition, 100 of the infants’ parents (74 mothers and 26 fathers) opted to contribute their own data to complement the data of their infant. Infants were in good general health, initially breastfed, and vaginally born between 37 and 43 weeks of gestation. Details about inclusion and exclusion criteria are provided in [33, 34, 35]. Ethical approval was obtained from the cantonal ethics committee (BASEC 2016-00730), and the study was carried out in accordance with the Declaration of Helsinki. Parents provided written informed consent. Families received small non-monetary gifts for their participation.

### Data collection

Activity data were recorded at 3, 6, and 12 months of age over 10 consecutive days each for a subset of 152 infant participants (n=150 at 3 months, 148 at 6 months, 143 at 12 months) and their parents (n=100 at 3 months, 79 at 6 months, 71 at 12 months), using actigraphy devices (GeneActiv, Activinsights Ltd, Kimbolton, UK). See Tables 1 and S1 for participant details. Parents were instructed to attach their own device to the non-dominant hand and the infant’s device to the ankle using a modified sock or a Tyvek paper strap. In addition, parents completed 24-hour diaries to document infant sleep episodes as well as times when the device was removed (primarily during bathing and swimming).

**Table 1.** Participant summary statistics. Number and frequency of participant characteristics in the infant and parental sample per time point. Infants were considered breastfed if breastfeeding frequency was reported as at least “4:regularly (3-5 times a week)”. Nighttime sleep indicates the relative amount of daily sleep occurring during nighttime per infant per time point. A family was counted as providing diadic or triadic data, whenever data from the infant plus one parent (diadic) or infant plus both parents (triadic) were available for the same time point.

|  |  |  | Time points |  |  |
| --- | --- | --- | --- | --- | --- |
|  |  |  | 3 months | 6 months | 12 months |
| Infants |  |  |  |  |  |
| Total |  |  | 150 | 148 | 143 |
| Females | n (%) |  | 67 (44.7%) | 67 (45.3%) | 64 (44.7%) |
| Breastfed | n (%) |  | 150 (100%) | 137 (92.6%) | 51 (36.4%)* |
| Sleep location |  |  |  |  |  |
| Separate room | n (%) |  | 20 (13.3%) | 41 (27.9%) | 83 (58.0%) |
| Separate bed | n (%) |  | 72 (48.0%) | 61 (41.5%) | 33 (23.1%) |
| Parents' bed | n (%) |  | 31 (20.7%) | 29 (19.7%) | 26 (18.2%) |
| Attached cot | n (%) |  | 27 (18.0%) | 16 (10.9%) | 1 (0.7%) |
| Nighttime sleep (%) | Median [IQR] |  | 72.3 [69.5, 72.5] | 78.6 [76.1,80.9] | 82.5 [80.1, 84.9] |
| Parents |  |  |  |  |  |
| Total |  |  | 100 | 79 | 71 |
| Mothers | n (%) |  | 74 (74%) | 61 (77.2%) | 56 (78.9%) |
| Families with diadic data | n |  | 54 | 45 | 40 |
| Families with triadic data | n |  | 23 | 17 | 15 |

\* out of 140

Feeding, sleeping habits as well as infants’ sleep location were also reported by the parents using questionnaires. At each time point, breastfeeding frequency was assessed on a 5-item scale, ranging from “1:never” to “3:occasionally (1-2 times a week)”, up to “4:regularly (3-5 times a week)” and “5:daily”. For the purposes of our analysis, breastfeeding was coded as “No” for levels 1 to 3 and as “Yes” for levels 4 and 5. Parents were also asked to describe their infant’s sleep location using a questionnaire providing 15 different sleep configurations (e.g.”Infant bed in separate room” or “In parents’ bed”). Here, these were classified with respect to their parents as “Separate room” or “Same room”. The latter category was further divided into “Separate bed”,”Same bed” or “Attached cot” (a semi-enclosed cot attached to the parental bed).

### Actigraphy processing

Actigraphy data pre-processing as well as sleep bout detection were performed with Matlab (R2016b), while all subsequent processing steps were performed in Python with the *pyActigraphy* software (v0.2) [36].

#### Pre-processing

As described elsewhere [33], raw accelerometry data acquired at a sampling frequency of 30 Hz were converted to activity counts in order to use validated sleep detection algorithms. In brief, following the method by Te Lindert et al. [37], a 3-11 Hz bandpass Butterworth (order 5) filter was applied to the magnitude of the acceleration vector along the 3 axes, which was then converted into counts by discretizing this value between 0 and 5g into 128 bins and summing the peak value measured every second over a period length of 1 minute.

#### Sleep bout detection and selection

Bouts of likely sleep (relative immobility), henceforth referred to as “sleep” for simplicity, were automatically identified from actigraphy recordings using the same procedure as in prior publications of this dataset, a modified version [33] of the algorithm developed by Sadeh and colleagues (1994) [38]. Modifications improved agreement between diary and actigraphy data and included an adapted threshold and a consolidation step where short active bouts (*≤* 5 min) surrounded by sleep were relabeled as sleep. Biases in recorded activity levels were reduced by excluding bouts where the infant was reportedly sick. Finally, from the sleep-wake scoring, the nighttime sleep percentage was calculated per infant per time point as the fraction of sleep duration occurring in the 12 hours between 19:00 and 07:00 with respect to the total sleep duration.

From all such estimated sleep bouts, we then selected only those occurring during the putative main sleep episode (defined as starting between 18:00 and 08:00) to minimize recording artifacts from external movement common during daytime at that age (from strollers, prams, carriers, etc.). We further selected bouts of a minimum duration of 90 min to enable adequate characterization of rhythms in the range of 40-90 min as informed by Singular Spectrum Analysis (see below). To prevent undue loss of data from this 90-min criterion and bias towards consolidated sleepers and later time points, we fused sleep bouts that were less than 15 min apart by padding the gaps with “Not a Number” (NaN) to represent missing data and retain the original temporal distance between the recorded activity patterns. Fused bouts were only included if they contained not more than 20% NaNs across their final duration. As a result, we were able to retain a median of 83 *−* 88% of sleep data per infant across all 3 time points (Table 2 and S2).

**Table 2.** Infant sleep bout summary statistics. Number and duration of actigraphically-identified nighttime sleep bouts of infants before and after processing and application of inclusion criteria. “Identified bouts” refers to the set of sleep bouts as detected by the modified Sadeh algorithm.”Analyzed bouts” refers to the final set of sleep bouts used in the inactivity analysis, after fusing of bouts within 15 minutes of each other and filtering for a 90-min minimum duration and a cosine fit offset outside of extreme values.

|  |  | Time points |  |  |
| --- | --- | --- | --- | --- |
|  |  | 3 months | 6 months | 12 months |
| Recording length (d) |  |  |  |  |
| Total/infant | Mean ± S.D | 11.1 ± 1.2 | 10.6 ± 1.9 | 10.6 ± 1.9 |
| Identified bouts |  |  |  |  |
| Sleep bouts (n) |  |  |  |  |
| Total |  | 8491 | 5600 | 3673 |
| Total/infant | Median [IQR] | 57 [46,68] | 38 [28,45] | 24 [19,31] |
| Total/infant/night | Median [IQR] | 5 [4,7] | 4 [3,5] | 2 [1,4] |
| Bout duration (h) |  |  |  |  |
| Total |  | 14698 | 13743 | 13565 |
| Total/infant/night | Median [IQR] | 9.9 [9.1,10.7] | 10.2 [9.4,10.9] | 10.6 [9.8,11.3] |
| Median/infant/night | Median [IQR] | 1.1 [0.8,1.9] | 2.1 [1.2,3.5] | 4.4 [2.0,6.1] |
| Analyzed bouts (fused if within 15 min; > 90 min; 1 <offset< 99) |  |  |  |  |
| Sleep bouts (n) |  |  |  |  |
| Total |  | 2479 | 1940 | 1524 |
| Total/infant | Median [IQR] | 17 [13,20] | 13 [10,15] | 10 [8,13] |
| Total/infant/night | Median [IQR] | 2 [1,2] | 1 [1,2] | 1 [1,2] |
| Bout duration (h) |  |  |  |  |
| Total |  | 11940 | 11978 | 11709 |
| Total/infant/night | Median [IQR] | 8.9 [7.1,10.1] | 9.8 [8.6,10.7] | 10.3 [9.4,11.1] |
| Median/infant/night | Median [IQR] | 4.8 [3.4,8.0] | 7.0 [4.6,10.0] | 9.9 [5.6,10.8] |

#### Locomotor Inactivity During Sleep (LIDS)

To study ultradian rhythms during sleep based on movement, activity counts during nighttime sleep bouts were converted to Locomotor Inactivity During Sleep (LIDS) following the procedure first laid out in Winnebeck et al. [20] with minor optimizing modifications. Formally, each detected sleep bout was processed as follows:

1. Non-linear transformation of activity to inactivity: LIDS_*i*_ = 100*/*(1 + *x*_*i*_), where *x*_*i*_ is the activity count at epoch *i*

2. Smoothing using a Gaussian kernel with a standard deviation (*σ*) of 5 min within a 30-min sliding window ([*−*3*σ*, +3*σ*])

An inactivity count of 100 represents an absence of movements, while the count tends to zero as the quantity (intensity and frequency) of movements increases. For ease of understanding, we refer to LIDS as “inactivity” throughout the manuscript outside of technical descriptions.

#### Singular Spectrum Analysis

To establish the presence of an oscillatory component in the infant inactivity signal, a Singular Spectrum Analysis (SSA) was performed. To avoid misinterpreting noise components as significant regular oscillations, a Monte Carlo SSA (MC-SSA) hypothesis test [39] from the python software *MCSSA* (v0.0.1) [40] was also used.

SSA aims to construct an additive decomposition of the original signal into a trend, regular oscillations, and a noise component [41]. Each component is associated with a singular value accounting for its partial variance. The MC-SSA then compares this signal decomposition to the decomposition obtained from an ensemble of random time series generated from a 1st-order autoregressive process (red noise). For each component, it is then possible to test the null hypothesis that this component originates from stochastic fluctuations.

To resolve infant periodicities in the same range as in our previous LIDS study in adults (up to 180 min), we chose an SSA embedding window length of 2.5 h. This meant that the required minimum sleep bout duration for SSA was 5 h, i.e. double the window length, as recommended for maximal separation of oscillations and noise [42]. The first detected sleep bout *>* 5 h from each infant at each time point was thus selected for SSA. Components with the 15 highest partial variances were kept, and components with a relative difference below 0.15 between their singular values were merged. These components were then statistically tested with MCSSA. Only statistically significant components were retained. Finally, to coarsely characterize each component, the cycle length associated with its maximal power, derived with a Fast Fourier Transform, was considered as the main cycle length.

#### Lomb-Scargle periodogram

In order to cross-check the presence of oscillations in infant inactivity, another periodic signal detection method was used: the Lomb-Scargle periodogram [43, 44] as implemented in the R package *MetaCycle* [45]. For each inactivity signal, the estimated cycle length corresponds to the highest periodogram peak; the null hypothesis that such a peak arose from white noise is then tested. A threshold at 0.05 on the statistical significance level was used to reject the null hypothesis and filter out statistically non-significant results. Two detection ranges (30-180 min, 15-240 min) were used to provide a coarse assessment of the robustness of the estimated periods.

#### Cosine curve fitting for LIDS parameter estimation

To quantify central features of the inactivity signal beyond cycle length (i.e. amplitude, phase, level at start and slope), a cosine model was fitted to the LIDS data using a non-linear leastsquares minimization from the package *lmfit* (1.0.3) [46]):

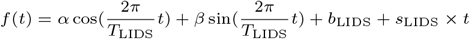

with

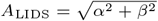: the LIDS amplitude,

*T*_LIDS_: the LIDS cycle length or period,

*ϕ*_LIDS_ = *−*1 *× atan*2(*β, α*): the LIDS phase at bout start (0 = peak at bout start; 180 = through at bout start),

*b*_LIDS_: the LIDS offset or vertical shift or inactivity level at bout start,

*s*_LIDS_: the LIDS slope or linear trend across the bout.

Similar to the approach in the original study, the fit procedure was performed iteratively; for each iteration, estimations of the model parameters were obtained by minimizing the sum of squared residuals, while the cycle length was fixed to a value ranging from 30-180 min in steps of 2 min. The selected best fit was the one corresponding to the peak value of the Munich Rhythmicity Index (MRI), defined as: MRI = 2 *× A*_LIDS_ *× r*, with *r*, the Pearson correlation coefficient between the inactivity data and the fitted model. Only sleep bouts with an offset greater than 1 and smaller than 99 were retained for statistical analyses to remove bouts with spurious activity patterns (e.g. no activity throughout the bout due to loss of the device during sleep).

#### Inactivity profiles

Plots of inactivity profiles were produced by averaging the inactivity signals of all relevant sleep bouts. Averaging was done based on three timelines: i) by absolute time since bout start, ii) by period-normalized time since bout start, iii) by period-normalized time since bout start after phase-alignment. Normalization and alignment reduce the destructive interference from intra- and inter-individual differences in LIDS cycle length and phase and thus better retain the oscillation in the averaged signal. Cycle length and phase were taken from the cosine model fits to each individual bout. The final visual display for the comparison between infants and their parents additionally rescaled the timeline to the median value of the cycle length extracted with the cosine model fitting, 62 min for infants and 80 min for parents.

#### Systematic uncertainties

In addition to reporting estimated mean parameter values and their statistical uncertainties (mainly arising from finite sample sizes), we also analyzed the systematic uncertainties arising from the methodological choices. Since the latter cannot simply be reduced by repeating the experiment and/or increasing sample size, their assessment is crucial to compare current results with those obtained with different methodologies from past and future studies. See S.I. for details and results.

### Statistical analysis

Generalized Bayesian linear mixed models, which account for the repeated measures nature of the data, were used to test for relationships between LIDS parameters (outcomes) and internal and external factors (predictors) such as time point, sex, sleep bout duration, percentage of nighttime sleep, breastfeeding, sleep location and/or family role.

First, for simple comparisons between infant and parent LIDS parameters at time point 1, models included a random intercept (*u*_*j*_) for each family (*j*) and status (infant or parent) as a predictor for each participant (*i*):

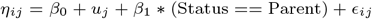

with *β*_0_ the estimated average LIDS parameter value for infants, *β*_1_ the estimated difference in the average LIDS parameter value between parents and infants, and *ϵ* the matrix of residuals.

Second, for more detailed analyses of LIDS parameters in infants including differences between time points, models included a random intercept (*u*_*i*_) for each participant (*i*) and a random slope (*u*_*ij*_) for each time point (*j*), with time point, sex and sleep location as categorical predictor variables and sleep bout duration as well as percentage of nighttime sleep as continuous predictor variables (fixed-effects). For these models, LIDS parameters extracted from all included sleep bouts were used as inputs. The general model structure was:

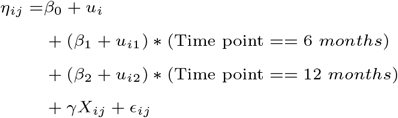

with *β*_0_ the estimated average LIDS parameter value, *β*_1_ and *β*_2_ the estimated differences in the average LIDS parameter value at 6 months (time point 2) and 12 months (time point 3) compared to 3 months (time point 1), *γ* the matrix of additional fixed-effects coefficients, *X* the matrix of their associated predictor variables, and *ϵ* the matrix of residuals.

The association between breastfeeding and LIDS parameters was also investigated with the above model, separately in infants and in mothers. This was done using data collected at 12 months only since there was little variation in breast-feeding frequency at the other time points. Hence, the model did not include time point as a predictor, but breastfeeding (yes/no) was added as a categorical predictor variable.

Third, a similar model was used to investigate the longitudinal modifications of LIDS parameters in parents. In this case, the model included a random intercept per family and random slopes for each time point. Time points, role (mother/-father), and sleep location were used as categorical predictor variables, and sleep bout duration as a continuous predictor variable. The model also included an interaction between role and time points.

Across all models, the linear predictor, *η*, was related to the outcome variable *y* (LIDS parameters) with the link function *g*:

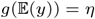

In parents, LIDS cycle length, amplitude, slope, and offset were modeled with a Student’s T distribution together with an identity link function. In infants, this was the same except for LIDS offset, for which a Gaussian distribution was the better fit. In all models for LIDS phase, the Von Mises circular distribution was used with a classic arctan link function.

A Bayesian mediation analysis [47] was performed to investigate whether increases in sleep bout duration indirectly mediated the LIDS cycle length increase with time. The model used for this analysis is identical to the default model used for the longitudinal analysis. However, for the sake of simplicity, time points were converted into months and used as a continuous independent variable.

All statistical analyses were performed in Python 3.8.10, using the Pandas (1.5.3) and Numpy (1.22.4) packages for data manipulation. Statistical Bayesian analyses were conducted with the Bambi (v0.10.0) software [48]. Unless stated otherwise, default weakly informative Gaussian priors were used. Posterior distributions were estimated with a Markov Chain Monte Carlo (MCMC) technique, sampling 4 independent chains in parallel, with 2500 draws for each chain. Convergence and overlap between chains were considered acceptable if the potential scale reduction factor 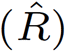 was smaller than 1.01. For each model parameter (*β*), the estimated posterior distribution was considered stable if its effective sample size was higher than 90% of the total number of draws (4 *×* 2500). Model adequacy was visually checked by computing the mean posterior predictive distribution. Finally, for each posterior distribution, we report its mean, standard deviation, and 95% highest density interval (HDI) as the credible interval (C.I.). Model parameter estimates were considered statistically significant if their C.I. did not overlap with zero.

## Results

Inactivity patterns in *>* 35, 000 hours of nighttime sleep were analyzed in detail in 152 infants during their first year of life, based on *∼* 10 days of ankle-actigraphy recordings at 3, 6, and 12 months of age. Almost all infants contributed to all 3 time points, and a consistent 45% were female (Tables 1 and 2). Additionally, in *>* 50% of families, at least one of the parents also contributed wrist-actigraphy recordings, providing a total of *>* 16, 000 hours of nighttime sleep recordings for parental inactivity pattern analyses. Mothers were the main contributors to the parental sample (74 *−* 79%). See Tables 1, S1 and S3 for more details on the parental sample and their sleep activity data contribution.

As expected the actigraphically-detected sleep-wake structure in infants changed between recording time points, becoming progressively less polyphasic and more consolidated. Specifically, sleep moved more into the night, increasing the proportion of daily sleep occurring at night from a median of 72.3% at 3 months to 82.5% at 12 months (Table 1), and was paralleled by an increase in the main nighttime sleep episode duration from a median of 9.9h per night per infant to 10.6h over the same time (Table 2). Therefore, we included an infant’s proportion of nighttime sleep in all key statistical analyses as a putative marker of maturation of the infant’s circadian and homeostatic system. Furthermore, nighttime sleep markedly consolidated over the assessment period. The median number of detected sleep bouts per night per infant evolved from 5 bouts at 3 months to 2 bouts at 12 months, with the median bout duration lengthening by 3.3 h, from 1.1 h to 4.4 h over this period (Table 2). Although we fused bouts separated by less than 15 min of wake for an optimal and minimally biased inactivity rhythm analysis (see Methods), sleep bout durations retained their characteristic variability within and between individuals and time points. We therefore also included bout duration as covariate in our statistical analyses. Importantly, the sleep-wake pattern development in our infant sample was aligned with nighttime sleep duration and consolidation reported in other infant cohorts (e.g. Rise&Shine cohort [49] and CHILD-SLEEP cohort [50]).

In addition to the longitudinal sleep-wake pattern changes, many of the parameters known or hypothesized to influence sleep quality, and thus considered as covariates in our analyses, also evolved markedly between the 3 recording time points (Table 1). While 100% of infants were breastfed at least 3 times a week at 3 months of age, this applied to only 36% at 12 months. Similarly, infants’ nighttime sleep location changed from predominantly the parents’ rooms and parents’ beds at 3 months (87%) to mostly separate rooms at 12 months (58%).

### Ultradian rhythms in infant inactivity during sleep

For an analysis of ultradian rhythmicity in infant movement during sleep, the activity signal of all sleep bouts was transformed to Locomotor Inactivity During Sleep (LIDS), enhancing the contrast between movement and non-movement (Figure 1A). This inactivity measure, ranging from 0-100 inactivity, formed the basis of all following analyses.

**Figure 1.**
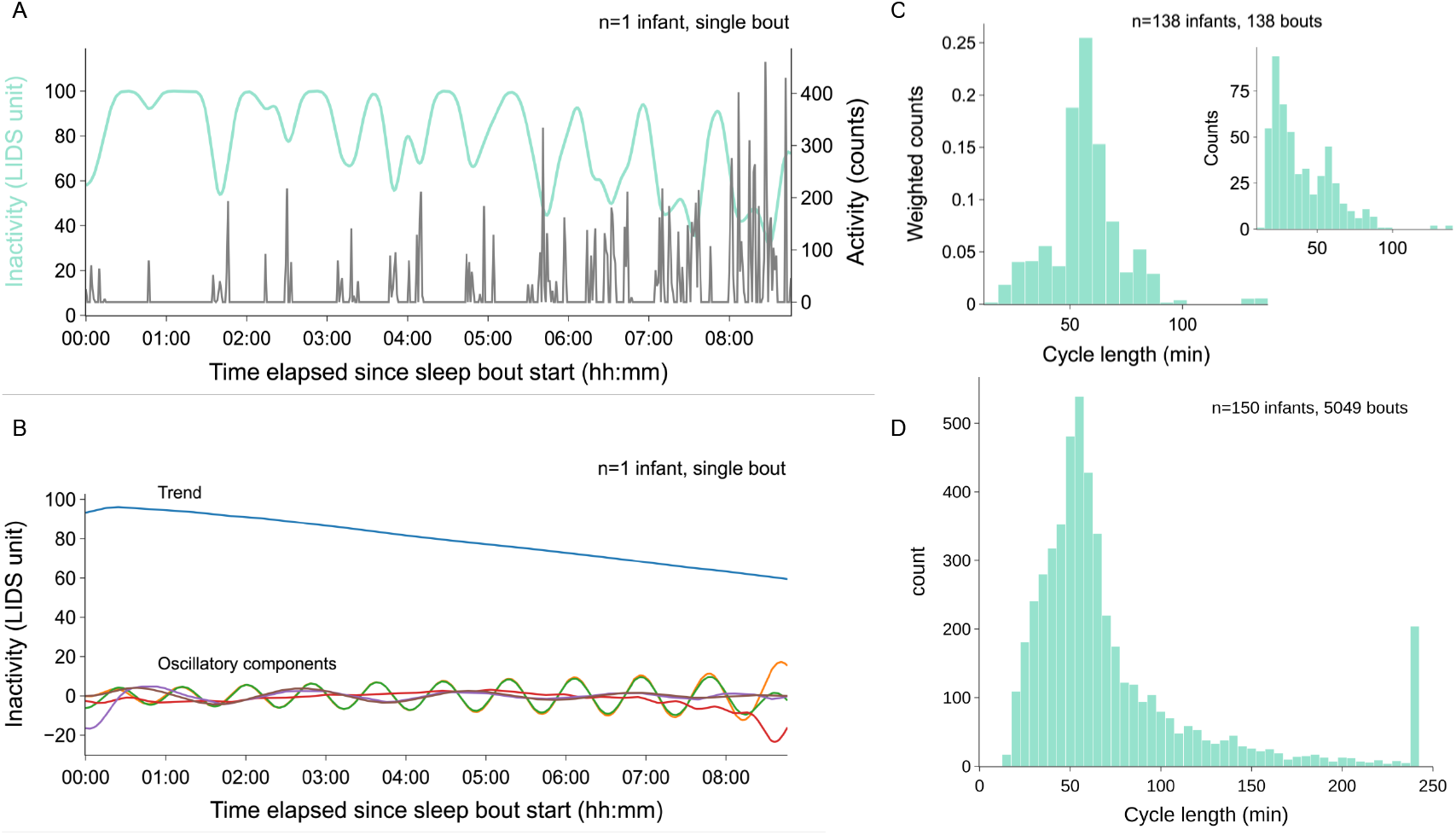
Locomotor Inactivity During Sleep (LIDS) in 3-month old infants oscillates with an ultradian cycle length of *∼* 60 minutes. **(A)** Example infant sleep bout showing locomotor activity (gray, right y-axis) and respective inactivity signal (turquoise, left y-axis). **(B)** Decomposition of the example inactivity signal from (A) into a trend and oscillatory components via Singular Spectrum Analysis (SSA). **(C)** Distributions of the main ultradian cycle lengths of statistically significant oscillatory components from SSA across infants’ first long sleep bout at 3 months. Histogram entries were weighted according to the amount of variance explained by their corresponding oscillatory component; inset displays the unweighted distribution. The analysis was based on the first bout *>* 5 hours for each infant at 3 months (1 bout per infant, n = 138 out of 150 infants, where sufficiently long bouts could be identified without fusing). **(D)** Distribution of the statistically significant cycle lengths from Lomb-Scargle periodogram analysis across all infant sleep bouts at 3 months.

#### Establishing rhythmicity

For a first ‘agnostic’ test of ultradian rhythmicity in infant inactivity during sleep, we decomposed the inactivity signal of a selected subsample of long bouts via Singular Spectrum Analysis (SSA). This separation of the signal into a trend and a set of oscillatory components (Figure 1B) revealed multiple statistically significant oscillatory components in infant inactivity already at 3 months. Figure 1C shows the distributions of the extracted main cycle lengths for each of these oscillatory components (often multiple per sleep bout). The unweighted histogram (inset) demonstrates that components with cycle lengths of 20-30 min were most abundant, followed by cycle lengths of 60 min. When weighted for the partial variance explained by each component, however, only the peak located around 60 min remained visible, indicating oscillations with a *∼* 60-min cycle length dominated the inactivity signals in infants at 3 months. This was confirmed by a second rhythm analysis, Lomb-Scargle periodograms across the full sample at 3 months, which also detected oscillations with a cycle length distribution peaking at *∼* 60 min (Figure 1D). Together, these results provide strong evidence for the existence of an ultradian rhythm in LIDS in 3-month-old infants with a main cycle length of around 60 min.

#### Rhythm characteristics at 3 months

The presence of such an ultradian inactivity rhythm as early as 3 months is evident also by visual inspection of the inactivity signal. Not only do most individual bouts show clear rhythms in inactivity (see example in Figure 1A), but also the averaged signal across all sleep bouts displays a rhythm in inactivity during infant sleep at all time points. While the oscillation of the averaged signal quickly dampens due to destructive interference from differing periods and phases (Figure S2A, B), period-normalization of the timeline and prior phase alignment of each bout’s inactivity signal reveals a strong, sustained oscillation in ankle inactivity during infant sleep (Figure 2A, B and Figure S2). Just like previously observed in adults, there is also a gradual decline in the levels of the inactivity oscillation over the course of sleep in infants.

**Figure 2.**
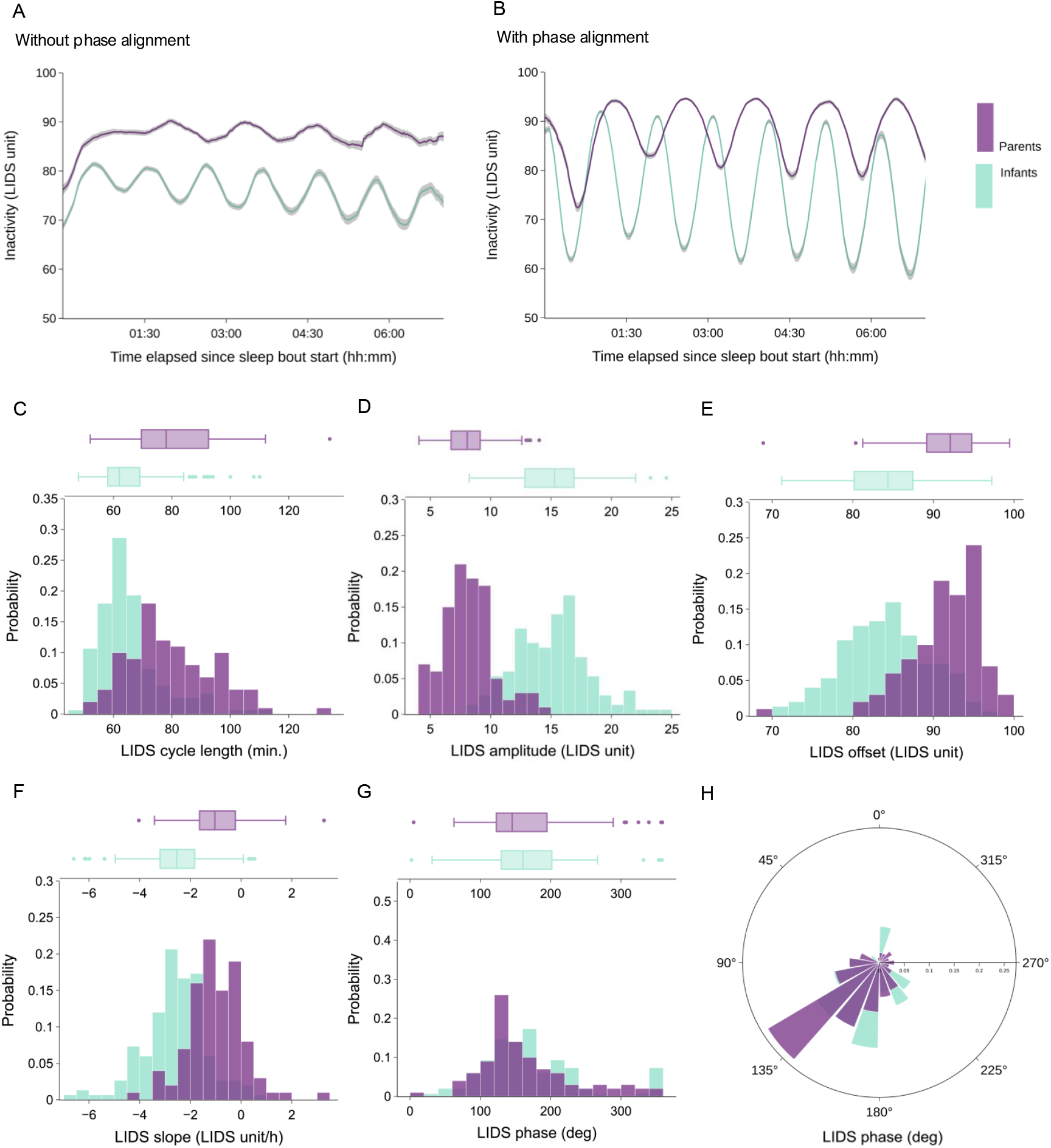
Comparison of inactivity rhythm parameters between infants and parents at 3 months. **(A**,**B)** Average inactivity profiles (*±*SEM) for infants and parents across all sleep bouts at 3 months. Signal timelines were normalized for inter-bout differences in cycle length before averaging and rescaled to infant and parent median LIDS cycle lengths for illustration, (A) without phase-alignment, (B) with phase-alignment. **(C-H)** Distributions of inactivity parameters for infants and parents at 3 months. Parameters were estimated via cosine fits to LIDS of individual sleep bouts and presented as medians per individual (C: cycle length, D: amplitude, E: offset, F: slope and G,H: phase). Boxplots are Tukey boxplots with whiskers spanning all data within 1.5 times the inter-quartile range above or below the central 50%.

To systematically characterize these ultradian inactivity rhythms in infants and compare them to those of their parents, parameter values were estimated via cosine model fits for LIDS cycle length, amplitude, offset, slope, and phase across the infant and parental samples. Below we consider these characteristics in detail at the 3-month time point, the first assessment time point in the sample, and thus earliest occurrence of the rhythm. Figure 2C-H shows the probability distributions of the estimated parameters at 3-months as medians per infant (turquoise) and per parent (purple). Most notably, the distribution of LIDS cycle lengths in infants exhibited a peak around 60 min (Figure 2C). This is in agreement with both the results from SSA in the subsample of long bouts (Figure 1C) and the results from Lomb-Scargle periodogram analysis in the full sample (Figure 1D). This 60-min cycle length also matches previous polysomnographically-determined cycle lengths of NREM-REM cycles in infants [29, 30], suggesting that infant inactivity oscillations might be similarly linked with sleep cycles as previously demonstrated in adults [20].

A further indication of such a link between inactivity and sleep cycles is the observation that the estimated LIDS cycle lengths in infants were shorter than those in adults, a well-established characteristic of polysomnographically-determined NREM-REM cycles. Importantly, at *∼* 60 min, LIDS cycle lengths were not only shorter than those previously reported in adults [20], but also shorter than those detected here in their parents (Figure 2A-C), which were recorded using the same device and the same sleep bout detection algorithm. Statistical testing using Bayesian generalized mixed model regressions indicated 12 min shorter median LIDS cycle lengths in infants than in their parents (Table S4).

Further visual and statistical comparisons between infants and parents at the 3-month time point revealed differences not only in LIDS cycle length but also in all other LIDS parameters except for LIDS phase (Figure 2 and Table S4). Parents had a smaller LIDS amplitude (*−*7.2 LIDS units), indicating that the oscillatory range in infants was larger than in parents, a higher LIDS offset (+7.4 LIDS units), indicating that infant inactivity was lower at the beginning of sleep than in parents, and a flatter LIDS slope (+1.6 LIDS units/h), indicating that the decline of inactivity over the course of sleep was steeper in infants than in parents. These results are in general agreement with results from the original LIDS analysis in children and adults [20], where LIDS cycle length was found to lengthen with age, from 5 to 92 years, while amplitude and decline were reduced. The results for phase do not provide evidence that infant and parent LIDS rhythms start at different points of the oscillation in relation to the actigraphy-defined sleep bout onset. With the modified Sadeh algorithm employed here, the inactivity rhythm was on average just before the trough at detected sleep onset, which is slightly earlier than with the MASDA algorithm in Winnebeck et al. [20], where it was around the peak.

Taken together, these observations suggest that infant inactivity during sleep is rhythmic, shows a cycle length of around 60 min, and differs systematically from parental inactivity patterns during sleep in multiple parameters.

### Factors associated with infant rhythm characteristics

#### Development: cycle length

To test for a lengthening of ultradian sleep cycles during infancy, we analyzed the evolution of inactivity cycle length in infants across the three assessment time points: 3, 6, and 12 months. As illustrated in Figure 3A, all three methods used for LIDS cycle length determination, SSA, Lomb-Scargle periodogram and cosine fits, indicate a gradual lengthening of inactivity cycles over the first year of life. The statistical analysis (Tables S5 and S6), based on the cosine fit estimates, puts the covariate-adjusted increase in LIDS cycle length at +4.7 min between 3 months and 12 months of age, while not detecting a statistically significant difference between 3 months and 6 months.

**Figure 3.**
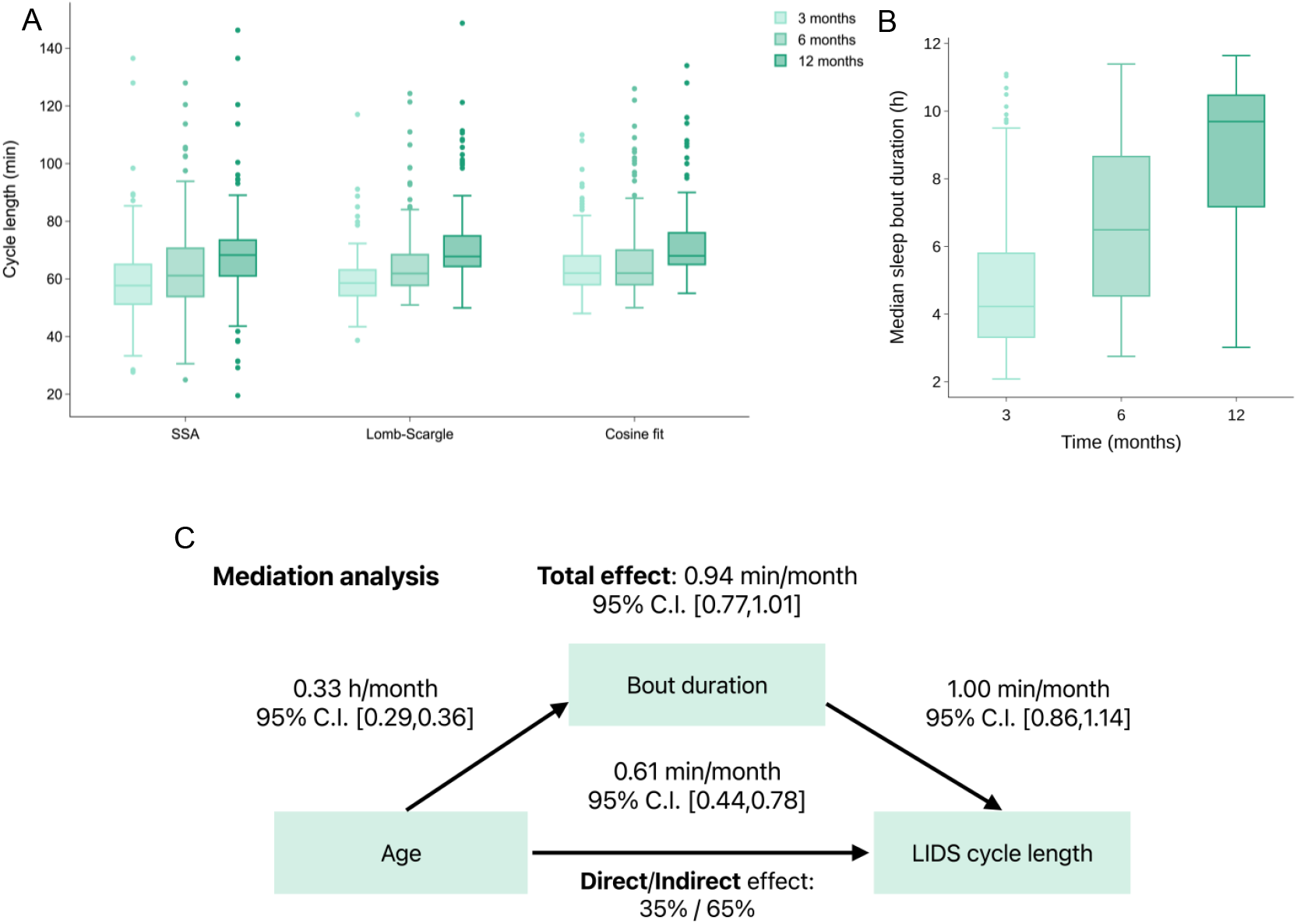
Infant inactivity rhythms lengthened over the first year of life. **(A)** Infant LIDS cycle length estimates at 3, 6 and 12 months obtained via 3 different methods: Singular Spectrum Analysis (SSA), Lomb-Scargle Periodogram (Lomb-Scargle) and Cosine fits. Note that samples differed between methods: SSA was run on 1 sleep bout per infant per time point (first bout *>* 5 hours), while the other 2 methods were applied to the full bout sample for which medians per infant are presented, explaining the lower variance in estimates compared to SSA. **(B)** Median sleep bout duration at 3, 6 and 12 months. **(C)** Schematic diagram of the mediation model to test the hypothesis that sleep bout duration mediated the increase in LIDS cycle length over time. Numerical values represent the corresponding fixed-effect coefficients and their 95 % credible intervals (C.I.). Boxplots are Tukey boxplots with whiskers spanning all data within 1.5 times the inter-quartile range above and below the central 50%. Please note that (A) shows unadjusted data and averages per individual. For more informative numbers, please refer to the statistically adjusted estimates reported in the text and tables.

Importantly, inactivity cycle length was also found to increase with sleep bout duration (+1.0 min/h, Table S5), an effect previously also detected in adults [20]. Since infant sleep bout duration increased markedly with age, doubling from 4.8h at 3 months to 9.9h at 12 months (Figure 3B; Table 2), the overall increase in LIDS cycle length between 3 and 12 months amounted to 9.8 min (4.7 min + 1.0 min/h *×* 5.1h). As expected, a similar increase was also found in a statistical model without adjusting for bout duration, which stipulates it at +7.6 min (Table S7).

Comparing the estimates between the models with and without adjustment for bout duration suggests that around a third of the increase of inactivity cycle length during infancy resulted from sleep bout lengthening, while two thirds arose independently. This indirect quantification is confirmed by a dedicated mediation analysis, which showed 35% of the effect of age on LIDS cycle length was mediated by the increase in bout duration, whereas 65% occurred independently (Fig 3C). Likewise, when selecting only bouts *>* 5 hours and restricting the cosine fit to the first 3 h of each bout, LIDS cycle length was still found to be longer at 12 months (+3.1 min) compared to 3 months (Table S8), demonstrating that cycle lengthening with age was detectable also in a sample of homogeneous bout durations and identical cosine fit lengths. Together, these results provide strong evidence for a gradual lengthening of ultradian sleep cycles during infancy and suggest potential concomitant lengthening mechanisms.

#### Development: other inactivity characteristics

Besides inactivity cycle length evolving over time in infants, all other LIDS characteristics, except phase, also changed throughout their first year of life (Figure 4). Compared to 3-month-old infants (Table S6), infants at 6 and 12 months had a lower LIDS amplitude (*−*2.3 LIDS units at 6 months, *−*1.6 LIDS units at 12 months). At the same time, LIDS offset increased (+3.3, +1.5 LIDS units) as did LIDS slope (+0.7, +0.8 LIDS units/h). The increase over time of LIDS offset shows that LIDS oscillations were initiated with a higher level of inactivity as the infants grew older, while the increase in the generally negative LIDS slope indicates a shallower decline in inactivity over the course of sleep at later time points. Together, this means that the overall level of inactivity remained higher throughout the sleep bout for older infants. As noted in [20], the probability of awakening most likely increases as the level of inactivity decreases. These results are thus compatible with infants having more consolidated sleep bouts and, therefore, longer sleep bouts during the night at 6 and 12 months compared to 3 months (Table 2).

**Figure 4.**
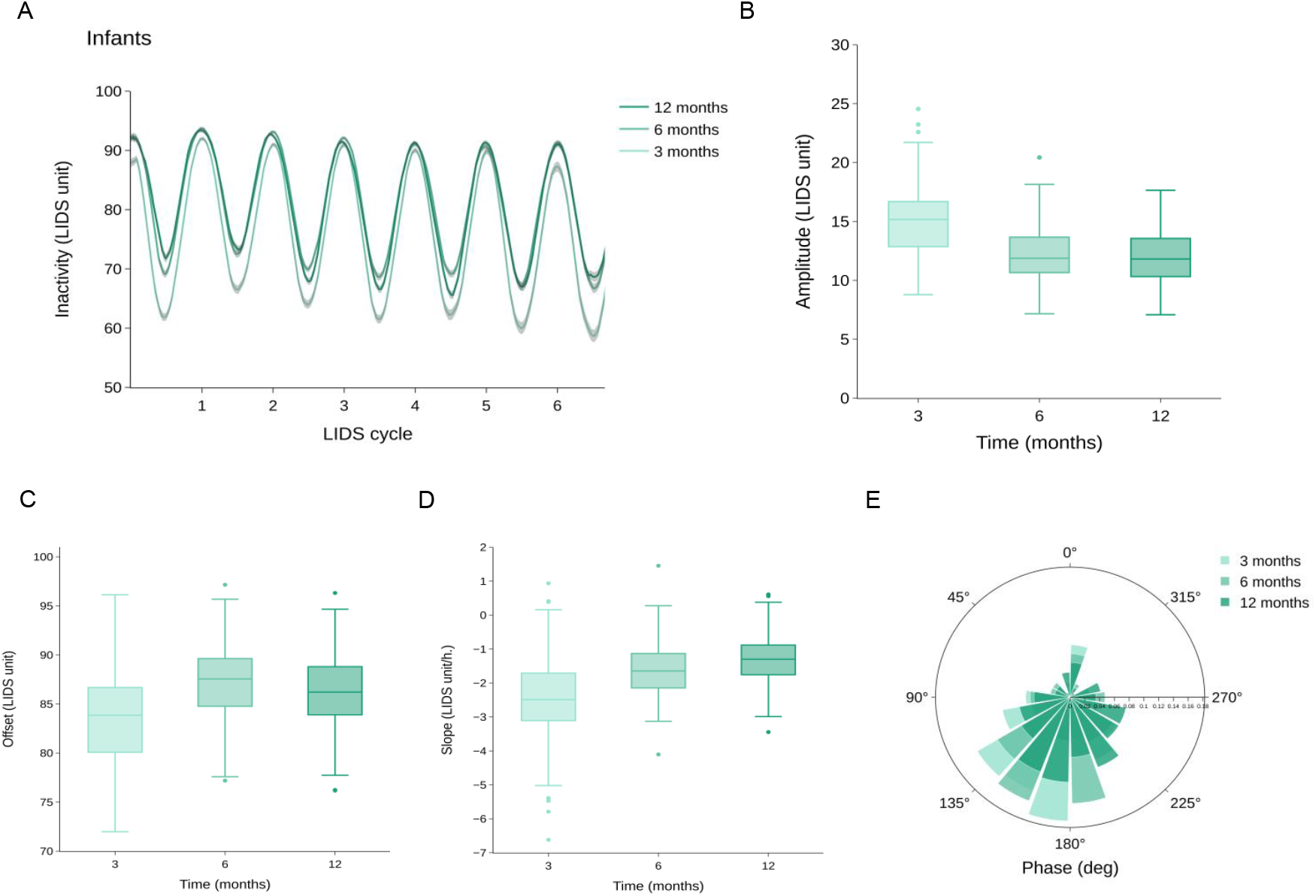
Development of other infant LIDS parameters over the first year of life. **(A)** Average inactivity profiles (*±*SEM) for infants at 3, 6 and 12 months. Signals were period-normalized and phase-aligned before averaging. **(B-E)** LIDS parameter estimates for infants at 3, 6 and 12 months. Parameters were estimated via cosine fits to LIDS of individual sleep bouts and are presented as medians per individual (B: amplitude, C: offset, D: slope, E: phase). Boxplots are Tukey boxplots with whiskers spanning all data within 1.5 times the inter-quartile range above or below the central 50%. Please note that the figures show unadjusted data and averages per individual. For more informative numbers, please refer to the statistically adjusted estimates reported in the text and tables.

No systematic differences in LIDS phase between time points were observed. However, it can be seen in the inactivity profiles (Figure S2) that, while rescaling the bout timelines with their individual LIDS cycle length seemed sufficient to reveal clear oscillations at 3 months, this was not the case at 6 and 12 months. This might point towards an increased variability of the LIDS phase at these time points, at least between individuals. Accordingly, aligning the LIDS phase between sleep bouts did restore visible oscillations at 6 and 12 months.

#### Endogenous factors: sex and sleep-wake cycle maturation

In none of the statistical models did the proportion of nighttime sleep show any association with any inactivity parameters. This is likely due to the low variability in this marker between infants and possibly its colinearity with bout duration and age.

Sex also explained little of the variation in infant inactivity parameters, with none of the effects reaching statistical significance. However, two patterns were notable: firstly, male infants tended towards longer sleep cycles (+1.3 min, 95% C.I.: [*−*0.08, +2.83]) across the full sample when accounting for sleep bout duration (Table S5). Secondly, LIDS offset tended towards lower levels in male infants (*−*0.8 LIDS units, [*−*1.77, +0.21]; Tab S6) suggesting greater movement during sleep in males. This matches several previous reports on sex differences in infant sleep motility, despite a lack of consistency across studies [51], as well as LIDS results in adults [20] and our result here in fathers (see below).

#### Exogenous factors: breastfeeding and sleep location

At 3 months, all infants in this study were regularly breastfed, while this was the case for just more than a third of the cohort at 12 months. We therefore explored differences in LIDS characteristics linked to differences in breastfeeding status at this last recording time point via a dedicated statistical model including breastfeeding status (Table S9). At 12 months, infants that were still breastfed at least 3-5 times per week had a longer LIDS cycle length (+2.5 min), as well as a higher LIDS offset (+1.4 LIDS unit), meaning that breastfed infants had longer inactivity cycles and initiated their sleep at a higher inactivity level compared to non-breastfed infants of the same age. Other LIDS parameters were not statistically linked to breastfeeding status.

In addition, we found associations between infants’ nighttime sleep location and LIDS characteristics (Tables S5 and S6). Compared to infants sleeping in a separate room, infants sleeping in their parents’ bed had longer LIDS periods (+2.9 min), lower LIDS amplitudes (*−*0.9 LIDS units) and a flatter LIDS decline (+0.3 LIDS units/h). These results were qualitatively identical to those obtained with the model including breastfeeding using sleep bouts collected at 12 months only (Table S9). Interestingly, the direction of the effects (increase or decrease) of sleeping predominantly in the parental bed and LIDS characteristics was similar to the effect of age.

### Factors associated with parental rhythm characteristics

#### Parental role

In parents, LIDS cycle length was stable across all time points. However, compared to mothers (Tables S10 and S11), fathers exhibited a shorter LIDS cycle length (*−*4.7 min) as well as a lower LIDS amplitude (*−*0.5 LIDS units).

Furthermore, fathers seemed to initiate their sleep at lower inactivity levels as reflected by a lower LIDS offset (*−*1.2 LIDS units), with the difference even further increased at 6 months by an additional *−*1.5 LIDS units. As the level of activity/inactivity in adults has been correlated with sleep depth [20, 52, 53], these results might be interpreted as deeper sleep/higher sleep pressure in mothers due to partial sleep deprivation, as expected for parents with young infants [54]. The fact that LIDS offset was also lower at 12 months, compared to 3 months, corroborates this interpretation.

Other LIDS parameters were not statistically different between mothers and fathers.

#### Breastfeeding (mothers only)

As breastfeeding might induce physiological changes that can, in turn, alter sleep in women, potential links between LIDS characteristics and maintained breastfeeding at the 12-months time point were also statistically assessed (Table S12). Indeed, at 12 months, compared to non-breastfeeding mothers, breastfeeding mothers had a longer LIDS cycle length (+6.7 min) as well as a steeper LIDS slope (*−*0.5 LIDS unit), indicating a faster decline of the inactivity levels. Other LIDS parameters were not statistically linked to breastfeeding habits.

It is interesting to note that both breastfed infants and breastfeeding mothers had a longer LIDS cycle length compared to their non-breastfeeding/fed counterparts.

#### Sleep location

The same statistical model for mothers at 12-months indicates associations between the sleep location of the infant and the mother’s LIDS characteristics (Table S12). Compared to mothers with an infant sleeping in a separate room, mothers sharing their room but not necessarily their bed with their infant had a longer LIDS cycle length (+8.6 min), and mothers sharing a bed with their infant had a greater LIDS amplitude (+1.8 LIDS units) and a lower LIDS offset (*−*2.8 LIDS units). Other LIDS parameters were not statistically linked to the mother-infant sleep settings.

Similar associations were found in the full parental model incorporating all time points and also the fathers (Tables S10 and S11), although the smaller sample size of fathers still biases this analysis towards the mothers. When families reported that their infant was sleeping predominantly in the parental bed, parents’ LIDS amplitude was again greater (+1.4 LIDS units) and LIDS offset was lower (*−*1.6 LIDS units) than when infants slept in a separate room. In contrast, in this model, the longest ultradian cycle lengths for parents were found when the infant slept in an attached cot (+6.3min), not in a separate bed as at 12 months.

## Discussion

Our study analyzed limb movement patterns during sleep as a window into infant and parental sleep physiology during infants’ first year of life. We extracted both rhythmic and non-rhythmic patterns of Locomotor Inactivity During Sleep (LIDS) from actigraphy and investigated associations with key endogenous and exogenous factors such as sex, sleep bout duration, breastfeeding, and sleep location to explore the potential of this approach for studying the functional relevance of sleep cycle dynamics in health and disease. Most notably, employing a range of careful analytic techniques, we demonstrated the existence of ultradian rhythms in infant inactivity that are compatible with inactivity patterns reflecting ultradian sleep cycles. These cycles lengthened between 3 and 12 months of age in concert with age and also with consolidation of sleep bouts, providing strong longitudinal evidence that sleep cycles lengthen gradually during development. Below we discuss the interpretation of the different inactivity pattern characteristics extracted from the LIDS signal, separating them by rhythmic and non-rhythmic components and by patterns associated with endogenous and exogenous factors.

A rich signal, such as limb movement during sleep, even with enhanced contrast between movement and non-movement via transformation to inactivity, inherently possesses many quantifiable features. Characterizing the LIDS signal using a cosine fit with a linear slope, we extracted five parameters of the inactivity signal during sleep. LIDS cycle length, amplitude, and phase are quantifications of the rhythmic aspects of the signal and might be interpreted as reflecting the periodicity, amplitude and phase of an underlying ultradian process. Phase, which quantifies the state of the inactivity rhythm at the beginning of the sleep bout, needs to be carefully interpreted as it is greatly dependent on the sleep-wake detection algorithm in cases where this is also based on movement like in our sample. In contrast, LIDS offset and LIDS slope quantify non-rhythmic aspects of inactivity during sleep, indicating the density and intensity of movement at the beginning of the bout (offset) and the gradual decline in inactivity with time asleep (slope). The latter reflects the previously observed gradual increase in movement during infant sleep [55]. Notably, amplitude also carries information about the amount of movement, and its extent is directly influenced by the intensity of movement through the non-linearity in the LIDS transform, therefore making amplitude a rhythmic and non-rhythmic parameter at the same time.

Based on prior research into body movements and sleep, including our own work on LIDS, [56, 52, 20], the most parsimonious interpretation of the non-rhythmic components extracted from LIDS is that they may reflect sleep depth and sleep pressure. Notably, deep sleep is more prominent in the first half of the night and is linked with lower motor activity. Thus, higher inactivity levels towards the beginning of a sleep bout likely correspond with deeper sleep early on in the bout [52, 53]. Another key feature of sleep is sleep homeostasis: the accumulation of sleep pressure during wakefulness and its gradual dissipation during subsequent sleep, in parallel with the decrease of deep sleep/slow wave activity over the course of the sleep bout [24, 57]. In infants, similarly to adults, mean inactivity levels decreased overnight in line with the gradual change from deeper sleep to lighter sleep [20]. Thus, the non-rhythmic components extracted from the LIDS signal may reflect both sleep depth and the dissipation of sleep pressure, an interpretation we use below for the findings in this study. However, it is without question that formal testing of this interpretation, using studies of experimental sleep restriction, will be essential to validate it.

Turning now first to the rhythmic patterns in inactivity during sleep, the central observation of our study was that motor inactivity during sleep is rhythmic in infants at 3 months, with a cycle length of approximately 60 min. This was established using three different analysis methods, Singular Spectrum Analysis, Lomb-Scargle periodogram and cosine model fitting. The results were not only in agreement with each other but also with the expected length of the NREM-REM sleep cycle for infants of this age [29, 30, 28]. Moreover, these findings go full circle with the original discovery of sleep cycles, where motility cycles were one of the main features characterized by Denisova, Aserinsky and Kleitman [14, 58, 13]. Additionally, infants showed shorter cycle lengths in inactivity than their parents, aligning with previous findings of different NREM-REM cycle lengths in infants and adults. This leads us to believe that both NREM-REM cycles and LIDS cycles are the manifestation of the same underlying ultradian process.

Furthermore, we demonstrated that inactivity cycles lengthen between 3 and 12 months of age by +4.7 min explained by the the main effect of age and by an additional +5.1 min explained by the increase in sleep duration. The developmental lengthening of LIDS cycles seems to parallel quite precisely the previously reported lengthening of NREM-REM sleep cycles in infancy [29, 30, 31, 28]. Indeed, in their longitudinal study of 15 infants, Louis et al. [30] demonstrated an increase in sleep cycle length of +11 min between 3 and 12 months, matching the increase we observed here (+9.8 min) over the same developmental period. Besides, in our study, this lengthening was evident across three different analysis techniques (SSA, Lomb-Scargle and cosine fit), which each have their own advantages and disadvantages in accommodating non-stationary signals, focus on cycle length determination or stability. Interestingly, the first two techniques, contrary to the cosine fits, indicated that lengthening occurred already between 3 and 6 months of age and therefore was gradual over the first year. This gradual nature of sleep cycle development in infants is corroborated by the cross-sectional report of Ficca et al. studying infants of diverse ages across the first year of life [31]. Our study adds substantial longitudinal evidence for gradual sleep cycle lengthening in a large sample of 150 infants over multiple consecutive nights per time point.

This central finding of developmental lengthening of ultradian sleep cycles could be linked to various phenomena, such as the maturation of an ultradian process underlying both NREM-REM and LIDS rhythmicity. As also observed in our sample (Table 2), the circadian and homeostatic systems mature with age [50, 49], resulting in more consolidated sleep bouts at night. Our mediation analysis revealed that longer sleep bout duration due to older age coincides with longer ultradian cycle lengths. Inter-individual differences in this system’s maturation may result in variability in the infant’s ability to generate consolidated and thus longer sleep bouts at night. But, as the cycles lengthen with age, longer cycles might also be considered as a sign of a more mature ultradian rhythm. Considered altogether, our observations of the endogenous factors affecting rhythmic components of sleep physiology suggest an interplay between the maturation of the ultradian and circadian systems.

When it comes to interpreting the exogenous effects on sleep cycles as gleaned from inactivity, the literature is more sparse, and our interpretations accordingly more speculative in nature. Nevertheless, we observed that breastfed infants at 12 months had longer LIDS cycle lengths, and therefore potentially a more mature ultradian rhythm than non-breastfed infants similar to older age. In addition, human breast milk composition exhibits diurnal variation; it presents higher levels of cortisol during the day and conversely higher levels of melatonin during the night [59]. Breastfeeding constitutes a hormonal *zeitgeber* that may facilitate circadian entrainment in infants and contribute to the maturation of the circadian system, as evidenced by an earlier appearance of a circadian rhythm in core body temperature in breastfed compared to non-breastfed infants [60]. This observation aligns with our previously stated hypothesis that circadian and ultradian rhythms are, in part, interrelated. Another explanation is that breastfeeding might also promote microbial diversity [61] and thus interact with the sleep-gut axis [35]. Indeed, within the same infant sample, a more diverse gut microbiome was linked to reduced daytime sleep. Considering that the circadian system actively promotes wakefulness during the day, less daytime sleep in infants might be interpreted as a more mature circadian system, which would in turn affect ultradian outputs. Conversely, a homeostatic explanation could be that the slower accumulation of homeostatic sleep pressure during wakefulness would also lead to less daytime sleep. Nonetheless, this observation should guide future studies, as interventions could be designed around modifiable factors that alter sleep and/or the gut microbiome during early infancy. Interestingly, breastfeeding mothers also showed a longer LIDS cycle, and further investigation is needed to determine whether this may also be linked to hormonal or microbial changes.

Turning to the non-rhythmic features of the inactivity signal, we observed multiple noteworthy associations with endogenous factors also in these. Firstly, LIDS offset (inactivity level at sleep onset) was lower in infants than in parents. This can be explained by the generally higher activity level of infants during sleep compared to adults, which is well documented and would result in lower LIDS offsets [51]. We also observed an increase in LIDS offset with age. If interpreting in the context of sleep homeostasis, this could potentially be indicative of higher sleep pressure in older infants, in line with our previous observation about decreased daytime sleep and consolidated, longer sleep bouts during nighttime. Additionally, LIDS slope (inactivity decline over the course of the sleep bout) showed a steeper decline in infants than in parents, possibly indicating a faster dissipation of homeostatic sleep pressure in infants, which is in line with mathematical models of infant sleep [62] and also with higher occurrence of slow-wave sleep in infants, a hallmark of sleep pressure dissipation, during the night [63]. However, infants maintained higher inactivity levels through the night as they grew older. It remains unclear if this is a cause or a consequence of sustained sleep episodes in older infants. Finally, LIDS amplitude was higher in infants than in parents, but decreased as infants aged. This phenomenon likely also reflects generally higher activity levels in infants. It remains to be investigated whether the pronounced LIDS amplitude in early infancy reflects the distinct neurophysiological features of infant sleep, as may be revealed by EEG markers of active and quiet sleep.

Finally, we also observed exogenous effects on the nonrhythmic aspect of inactivity in what is a highly exploratory analysis of this topic, given the small prior number of studies on the subject. Breastfeeding mothers showed a steeper decline in inactivity over the course of sleep compared to non-breastfeeding mothers. This is interesting in light of the increased sleep pressure seen in breastfeeding mothers, as suggested by a higher delta power in their night sleep electroencephalogram (EEG) power spectrum and more time spent in deep sleep [64, 65]. This increase has been shown to remain in breastfeeding mothers even when controlling for partial sleep deprivation, suggesting that these differences in nighttime sleep EEG characteristics might be, at least partially, due to physiological adaptations related to increased circulating prolactin levels induced by breastfeeding. When it comes to sleep environmental influences, various sleep characteristics, such as sleep duration or quality, have been shown to be modulated by socioeconomic and environmental factors [66, 67]. Parent-infant sleep settings (solitary sleeping infants or infants sharing a room and/or bed with their parents) seem to affect both infants and mothers, as well as mothers’ postpartum sleep trajectory [68, 69]. Interestingly, our exploratory analysis revealed that LIDS characteristics were linked to bed/room sharing habits, both in infants and parents, suggesting that dedicated studies on the impact of the environment on sleep might benefit from the LIDS methodology.

## Limitations

While this study demonstrates the great potential of using large longitudinal activity records as a window into infant and parent sleep, we note several limitations for the interpretation of this study. It cannot be excluded that differences in inactivity patterns between infants and parents partially originated from the different placement of devices between the two groups. Actigraphy data were collected using ankle-worn devices in infants while parents wore the actigraph at their wrists. However, it was demonstrated that, at least in adults, ankle-derived and wrist-derived LIDS patterns are highly correlated within individuals and sleep bouts [20].Additionally, both infants and parents were recorded using the exact same device model, and the same sleep-wake algorithm was used on all recordings, limiting methodological differences from these aspects. Another limiting factor arose from the natural settings in which actigraphy data were collected; while measuring sleep in the habitual sleep environment underscores the study’s relevance, it also hampers exclusion of confounding factors, such as environmental noise. This is particularly relevant for our exploratory analyses of breast-feeding and sleep location, which need to be interpreted carefully and used only for hypothesis generation to be followed up in more controlled studies. Finally, the social, economic, and cultural characteristics of the participants should also be taken into account when generalizing the study findings, as they have been shown to exert an influence on sleep characteristics [66]. Reproducing our results with culturally and racially diverse cohorts is therefore mandatory.

## Conclusions

Epidemiological, longitudinal and experimental research has demonstrated the importance of sleep for infant brain development and cognitive functioning. Methods that allow large-scale characterization of sleep beyond simple sleep duration commonly obtained from self-reports or actigraphy will be essential in enabling better understanding of the interplay between sleep physiology and development. Even though movement records will unlikely ever convey as much detail as polysomnography, the simplicity and unobtrusiveness of data recording enables large, longitudinal samples that are instrumental for hypothesis generation, in discovering potential genetic mechanisms through genome-wide association studies or identifying potential biomarkers for developmental progress that are suitable as screening tools. Using limb movement from actigraphy or other wearables as a window into sleep physiology, and particularly the thus far still enigmatic ultradian sleep cycles, promises great potential in advancing our understanding of the intricate relationship between sleep and developmental trajectories.

## Acknowledgment

We would like to thank the families for their participation in this study.

## Author contributions statement

- Inactivity pattern analysis

– GH: Formal Analysis; Methodology; Software; Validation; Visualization; Writing – Original Draft Preparation; Writing – Review & Editing
– SFS: Data curation; Formal Analysis; Writing – Review & Editing
– ME: Formal Analysis; Validation
– ZS: Validation; Writing – Review & Editing
– SK: Writing – Review & Editing
– ECW: Conceptualization; Funding Acquisition; Methodology; Project Administration; Supervision; Validation; Writing - Original Draft Preparation; Writing – Review & Editing

- Infant cohort

– SFS: Data curation; Formal Analysis; Funding Acquisition; Investigation; Methodology; Project Administration
– SK: Conceptualization; Funding Acquisition; Investigation; Project Administration

## Supplementary information

### Systematic uncertainties on estimated cosine model parameters

Systematic uncertainties of the estimated infant cosine model parameter X, 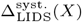, were assessed by computing the absolute difference between the median value per participant obtained with the default settings and the median value for the same participant with a modified setting. In this study, we investigated systematic uncertainties from the following alternative settings:

1. Inactivity signal processing:
  a. Smoothing: Inactivity data smoothed using a centered moving average with a window length of 30 minutes, as used in the original LIDS study.
  b. Smoothing kernel resolution: Inactivity data smoothed using a Gaussian kernel with a standard deviation ranging from 2.5 to 10 minutes.
  c. Sampling resolution: Inactivity data resampled to a 10-min resolution, the resolution in the original LIDS study. However, since smoothing data with 10-min wide bins with a Gaussian kernel with a 5-min resolution is inefficient, a centered moving average with a window length of 30 minutes was used instead, both for the original and the resampled data.
2. Linear trend in LIDS harmonic model: LIDS parameter estimated with a model without linear trend, as in the original LIDS study.
3. Sleep bout fusing: unfused sleep bouts as well as with a maximal time gap of 5 minutes.

#### LIDS cycle length

In infants, the source of systematic uncertainties on the LIDS cycle length estimates with the greatest magnitude is the smoothing procedure. LIDS smoothed with a 30-min long moving average window yielded longer LIDS cycle length estimates (+6 min (*IQR* = [+3, +12]) at 3 months, +6 min (*IQR* = [+2, +15]) at 6 months and +3 min (*IQR* = [+0, +10]) at 12 months). Similarly, a smoothing using a wider Gaussian kernel (*σ* = 10 min) led to longer LIDS cycle length estimates (+8 min (*IQR* = [+4, +16]) at 3 months, +10 min (*IQR* = [+4, +22]) at 6 months and +4 min (*IQR* = [+2, +17]) at 12 months). No sleep bout fusing or a fusing with a time gap of 5 minutes appeared to decrease the LIDS cycle length estimates (*−*2 min (*IQR* = [*−*6, +0]) at 3 months, *−*3 min (*IQR* = [*−*8, +0]) at 6 months and *−*2 min (*IQR* = [*−*6, +0]) at 12 months). This observation was compatible with the increase in LIDS bout duration (+61 min, *IQR* = [+20, +130]) with the bout fusing, which in turn produces longer LIDS cycle length estimates (cf. Table S5). A smoothing with a narrower Gaussian kernel (2.5 min), a lower sampling bin width (10 min) of the sleep bouts, as well as the addition of a linear trend in the LIDS oscillatory model had marginal effects on LIDS cycle length estimates.

#### Other LIDS characteristics

As expected, the main source of systematic uncertainty of the amplitude estimates originated from the sampling frequency: +11.1 LIDS unit (*IQR* = [+9.6, +12.8]) at 3 months, +10.7 LIDS unit (*IQR* = [+9.3, +12.4]) at 6 months and +10.8 LIDS unit (*IQR* = [+9.4, +12.6]) at 12 months with a lower resampling resolution (10 min), compared to estimates produced with the original frequency (1 min). Since the resampling was performed via summation over aggregated epochs, the mean LIDS levels, and therefore the LIDS amplitude, increased almost linearly with the resampling resolution. In addition, the smoothing resolution was the second largest source of systematic uncertainties for the LIDS amplitude. Compared to the default smoothing resolution, a lower temporal resolution yielded lower LIDS amplitude; a wider Gaussian kernel leads to *−*4.1 LIDS unit (*IQR* = [*−*5.1, *−*3.3]) at 3 months, *−*3.0 LIDS unit (*IQR* = [*−*3.9, *−*2.2]) at 6 months, *−*2.6 LIDS unit (*IQR* = [*−*3.3, *−*2.1]) at 12 months and a 30-min long moving average window to *−*3.5 LIDS unit (*IQR* = [*−*4.3, *−*2.6]) at 3 months, *−*2.5 LIDS unit (*IQR* = [*−*3.3, *−*1.8]) at 6 months and *−*2.1 LIDS unit (*IQR* = [*−*2.7, *−*1.6]) at 12 months. This effect was expected; the lower the temporal resolution (i.e. the larger the smoothing kernel), the higher the loss of temporal specificity. LIDS levels were spread over larger time periods and therefore decreased. This observation emphasized that the LIDS level fluctuations are temporally localized and not random as expected for rhythmic processes. Other sources of potential systematic uncertainties had marginal effects on the LIDS amplitude estimates.

Similarly to the LIDS amplitude, the LIDS offset was affected by the resampling frequency: *−*28.8 LIDS unit (*IQR* = [*−*33.5, *−*24.6]) at 3 months, *−*25.6 LIDS unit (*IQR* = [*−*29.4, *−*21.3]) at 6 months and *−*26.7 LIDS unit (*IQR* = [*−*29.7, *−*24.6]) at 12 months for a lower resampling frequency (1*/*10 min^*−*1^). Another prominent source of systematic uncertainty derived from the model parametrization; obviously, the addition of a linear trend in the LIDS model, compared to the original model, modified the LIDS characteristic estimated by the offset. In the current model, the offset represented the LIDS level at the beginning of the sleep bout while, in the original model, it represented the overall mean LIDS level. As the LIDS levels decreased overnight, the mean LIDS levels were lower than the LIDS levels at the beginning of the sleep bout: *−*7.1 LIDS unit (*IQR* = [*−*9.7, *−*4.5]) at 3 months, *−*6.1 LIDS unit (*IQR* = [*−*7.8, *−*4.0]) at 6 months, *−*5.4 LIDS unit (*IQR* = [*−*7.9, *−*3.4]) at 12 months. The decrease of this systematic effect over the time points (from *−*7.1 to *−*5.4 LIDS unit) was consistent with the concomitant increase of the LIDS slope (cf. Table S6).

### Summary statistics for participants and sleep bouts

Table S1 provides information about the number of participants as a function of their family members enrolled in this study. In addition, Table S3 describes the number of sleep bouts identified in parents, as well as their duration, across the three assessment time points. Similarly to the infants, the median number of sleep bouts per night decreased from 4 *−* 5 to 1 when fusing and cosine fit quality criteria were applied. In parallel, the median sleep bout duration per night increased from 0.8h to 5.8h at 3 months and from 0.9h to 6.9h at 12 months. This new duration is most likely more representative of the expected average sleep duration in young adults.

## Supplementary figures

Lomb-Scargle periodogram

**Figure S1.**
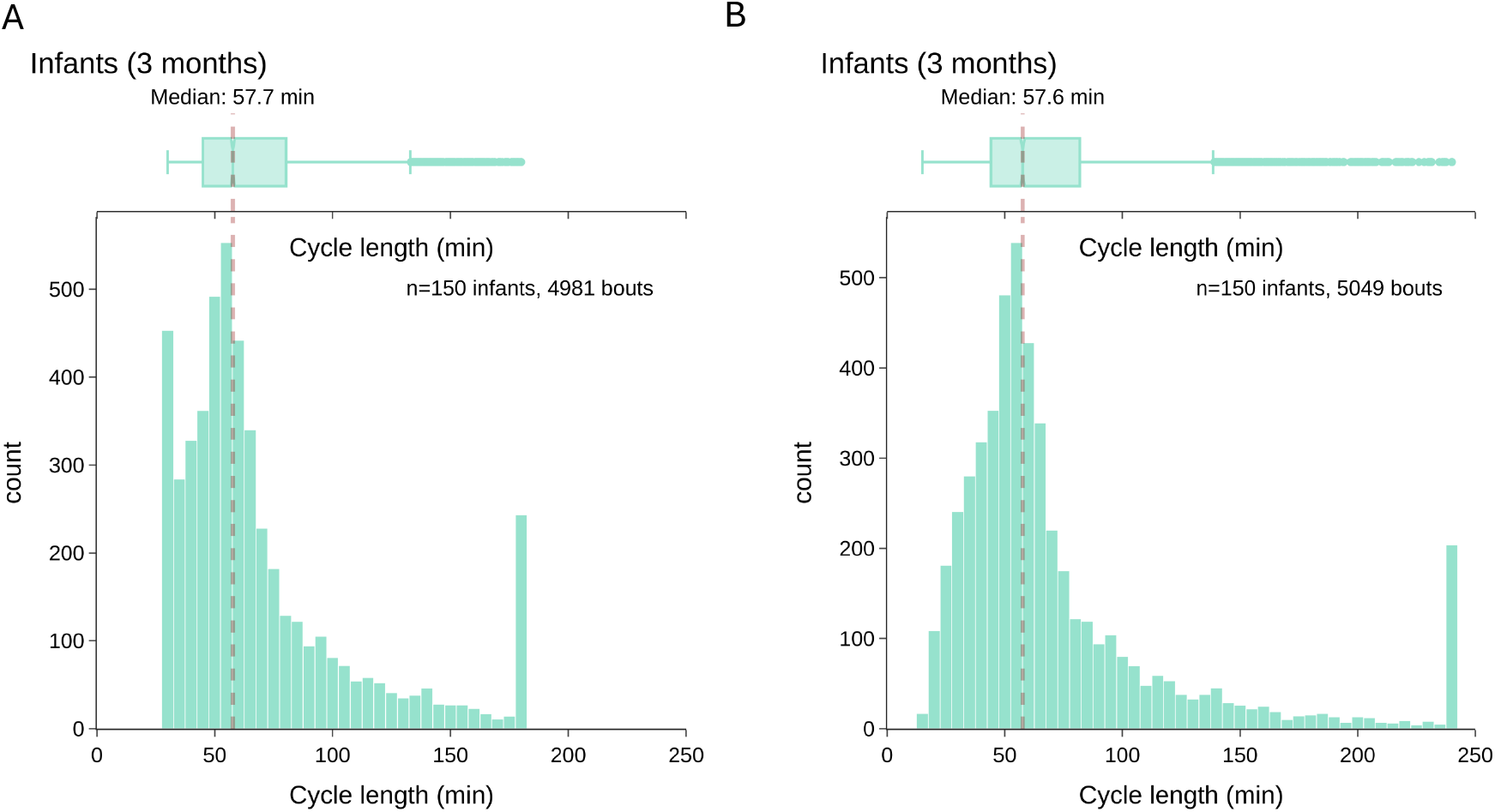
Lomb-Scargle determined cycle lengths of the LIDS rhythm in infants at 3 months. Distribution of estimated cycle length of the full infant sleep bout sample at 3 months as determined by Lomb-Scargle periodogram analysis with two different detection ranges: (A) 30-180 min and 15-240 min cycle lengths. Boxplots are Tukey boxplots with whiskers spanning all data within 1.5 times the inter-quartile range. Spikes at a cycle length of 30, 180 and 240 minutes are due to bouts with an estimated cycle length at the limits or outside of the detection range.

### Inactivity profiles

Figure S2 shows the infant inactivity profiles across the first year of life.

**Figure S2.**
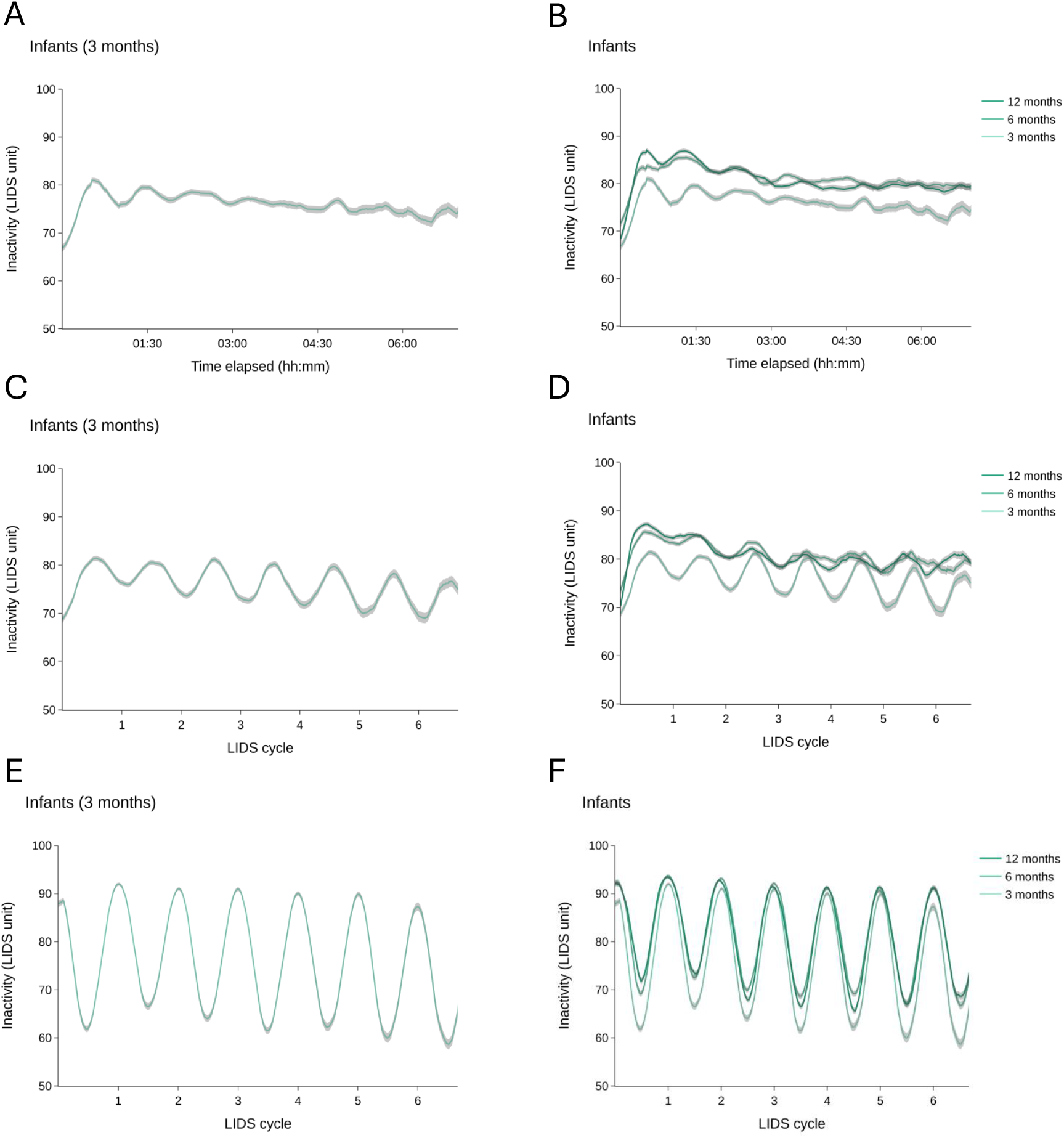
Infant inactivity profiles across the first year of life. Average profiles (*±*SEM) of infant inactivity during sleep at 3 months (left) and all 3 time points (right). **(A**,**B)** “Raw” profiles, where inactivity was averaged across sleep bouts by absolute time since bout onset. **(C**,**D)** Period-normalized profiles, where inactivity was averaged by time normalized to each bouts’ LIDS period. **(E**,**F)** Phase-aligned and period-normalized profiles, where inactivity was averaged by time normalized to each bout’s LIDS period after phase-alignment according to each bout’s LIDS phase.

### Systematic uncertainties on estimated cosine model parameters

**Figure S3.**
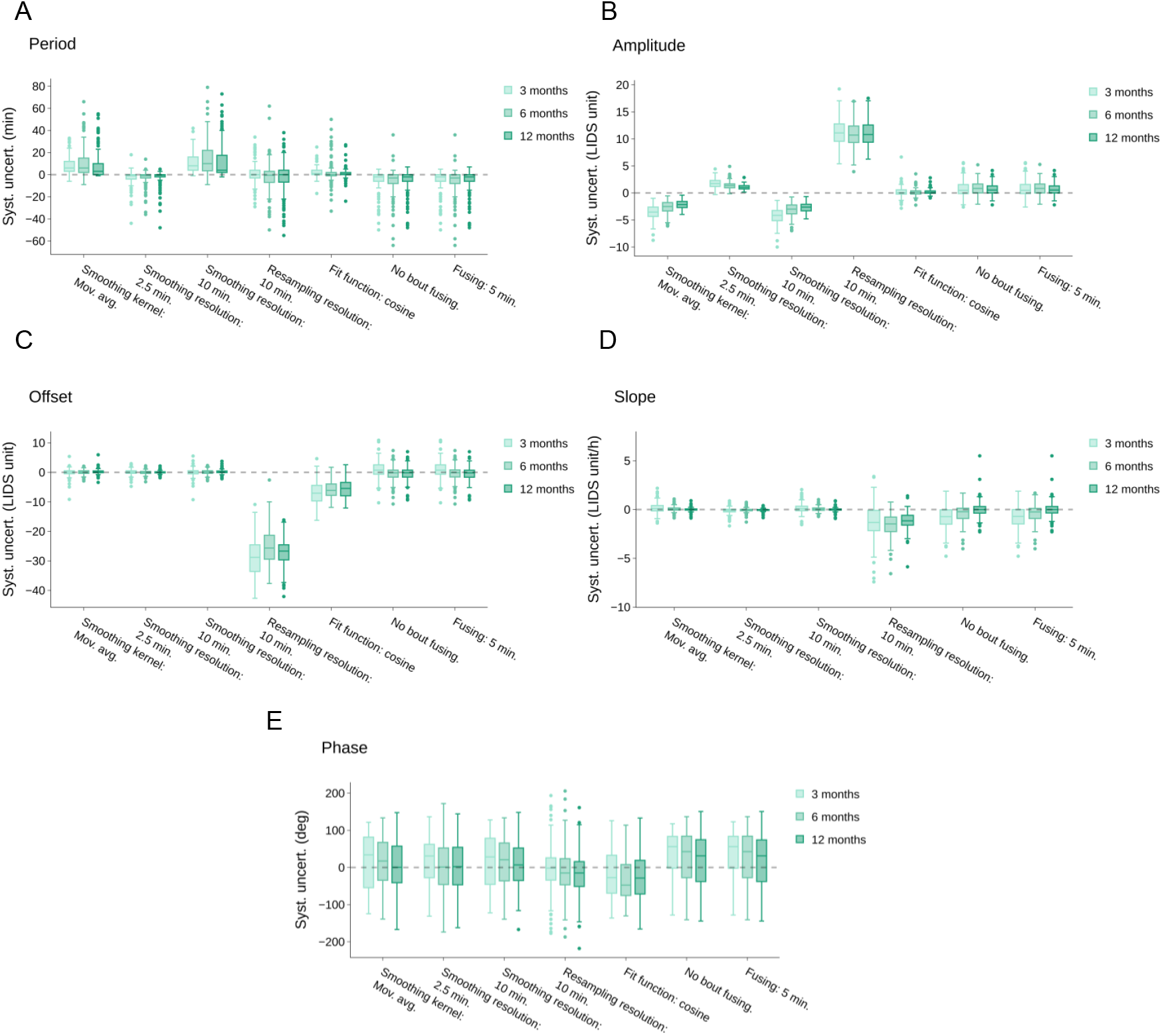
Systematic uncertainties of LIDS parameter estimation in infants. The systematic influence of key data processing choices on LIDS parameter estimation. Displayed are differences between the individual median values obtained with the default processing choice to the individual median values obtained with the altered settings indicated. All LIDS parameter estimates were performed via cosine fitting (A:period, B:amplitude, C:offset, D:slope and E:phase). Boxplots are Tukey boxplots with whiskers spanning all data within 1.5 times the inter-quartile range.

**Table S1.** Data availability per family. Number of families for which data were available for analysis as a function of contributing family members at time points 1 (3 months) to 3 (12 months). Abbreviations: I, infant; M, mother; F, father

| Family contribution | I only | M only | I + M | I + F | I + M + F | Total families<br>I + any parent | Total families<br>I only or I + any parent |
| --- | --- | --- | --- | --- | --- | --- | --- |
| Time point |  |  |  |  |  |  |  |
| 3 months | 73 | 0 | 51 | 3 | 23 | 77 | 150 |
| 6 months | 86 | 0 | 44 | 1 | 17 | 62 | 148 |
| 12 months | 88 | 1 | 40 | 0 | 15 | 55 | 144 |

## Supplementary tables

Summary statistics for participants and sleep bouts

**Table S2.** Infant sleep bout summary statistics. Number and duration of actigraphically-identified nighttime sleep bouts of infants after fusing of bouts within 15 minutes of each other (excluding those comprising *>* 20% wake from fusing).

|  |  | Time points |  |  |
| --- | --- | --- | --- | --- |
|  |  | 3 months | 6 months | 12 months |
| Sleep bouts after fusing (< 20% wake) |  |  |  |  |
| Sleep bouts (n) |  |  |  |  |
| Total |  | 3496 | 2408 | 1742 |
| Total/infant | Median [IQR] | 23 [19,27] | 17 [12,19.25] | 11 [9,15] |
| Total/infant/night | Median [IQR] | 2 [2,3] | 2 [1,2] | 1 [1,2] |
| Bout duration (h) |  |  |  |  |
| Total |  | 13423 | 12920 | 12137 |
| Total/infant | Median [IQR] | 92.3 [81.1,99.6] | 92.7 [80.2,101.0] | 90.0 [74.3,101.2] |
| Total/infant/night | Median [IQR] | 9.5 [8.3,10.5] | 10.1 [9.2,10.9] | 10.4 [9.6,11.1] |
| Median/infant/night | Median [IQR] | 4.0 [2.4,5.5] | 5.4 [4.2,9.9] | 9.7 [5.2,10.8] |

**Table S3.** Parent sleep bout summary statistics. Number and duration of actigraphically-identified nighttime sleep bouts of parents before and after processing and application of inclusion criteria. “Identified bouts” refers to sleep bouts as detected by the modified Sadeh algorithm. “Fused bouts” refers to sleep bouts available after concatenation of bouts within 15 minutes of each other (after exclusion of those comprising *>* 20% wake from fusing).”Analyzed bouts” refers to the final set of sleep bouts used in the inactivity analysis after filtering for a 90-min minimum duration and a cosine fit offset outside of extreme values. Abbreviation: IQR, interquartile range

|  |  | Time points |  |  |
| --- | --- | --- | --- | --- |
|  |  | 3 months | 6 months | 12 months |
| Identified bouts |  |  |  |  |
| Sleep bouts (n) |  |  |  |  |
| Total |  | 4933 | 3517 | 2738 |
| Total/parent/night | Median [IQR] | 5.0 [3.0,7.0] | 4.0 [3.0,6.0] | 4.0 [3.0,5.0] |
| Bout duration (h) |  |  |  |  |
| Total |  | 8088 | 6340 | 5256 |
| Total/parent | Median [IQR] | 83.3 [72.2,89.7] | 81.2 [73.8,87.3] | 77.2 [69.6,84.4] |
| Median/parent/night | Median [IQR] | 0.8 [0.4,1.7] | 1.0 [0.5,2.0] | 0.9 [0.5,2.5] |
| Fused bouts (fused if within 15 min; < 20% wake) |  |  |  |  |
| Sleep bouts (n) |  |  |  |  |
| Total |  | 1792 | 1288 | 1028 |
| Total/parent/night | Median [IQR] | 2.0 [1.0,2.0] | 2.0 [1.0,2.0] | 1.0 [1.0,2.0] |
| Bout duration (h) |  |  |  |  |
| Total |  | 7606 | 5984 | 5005 |
| Total/parent | Median [IQR] | 79.4 [68.4,85.8] | 77.6 [68.9,83.1] | 72.4 [66.2,80.3] |
| Median/parent/night | Median [IQR] | 4.3 [3.0,7.4] | 4.8 [3.5,7.7] | 5.4 [3.7,7.7] |
| Analyzed bouts (fused if within 15 min; > 90 min; 1 <offset< 99) |  |  |  |  |
| Sleep bouts (n) |  |  |  |  |
| Total |  | 1226 | 935 | 747 |
| Total/parent/night | Median [IQR] | 1 [1,2] | 1 [1,2] | 1 [1,1] |
| Bout duration (h) |  |  |  |  |
| Total |  | 6490 | 5353 | 4454 |
| Total/parent | Median [IQR] | 66.6 [53.4,77.2] | 69.9 [56.5,77.3] | 63.9 [57.7,71.2] |
| Median/parent/night | Median [IQR] | 5.8 [3.8,8.1] | 6.7 [4.2,8.2] | 6.9 [4.8,8.0] |

Statistical results

**Table S4.** Statistical comparison of LIDS parameters between infants and parents at 3 months. Bayesian generalized mixed model analysis with random intercept per family: estimated mean, standard deviation and limits of the credible interval of posterior distributions for fixed and random effects. Statistically significant effects are shaded in gray. Reference levels for categorical fixed effects: Infant for Role. Abbreviations: Std, standard deviation and HDI, highest density interval.

| LIDS parameter |  | Mean | Std. | 95% HDI:<br>lower limit | 95% HDI:<br>upper limit |
| --- | --- | --- | --- | --- | --- |
| Cycle length (min) | Intercept | 66.17 | 0.79 | 64.60 | 67.70 |
|  | Role[Parent] | 11.75 | 1.01 | 9.82 | 13.77 |
| | Random effect: $\sigma_1\vert _{\text{Family ID}}$ | 5.87 | 0.64 | 4.67 | 7.15 |
| Amplitude (LIDS unit) | Intercept | 15.18 | 0.20 | 14.79 | 15.56 |
|  | Role[Parent] | -7.17 | 0.17 | -7.49 | -6.83 |
| | Random effect: $\sigma_1\vert _{\text{Family ID}}$ | 2.13 | 0.15 | 1.83 | 2.43 |
| Offset (LIDS unit) | Intercept | 84.45 | 0.35 | 83.77 | 85.13 |
|  | Role[Parent] | 7.42 | 0.36 | 6.69 | 8.11 |
| | Random effect: $\sigma_1\vert _{\text{Family ID}}$ | 3.31 | 0.28 | 2.78 | 3.86 |
| Slope (LIDS unit/h) | Intercept | -2.56 | 0.08 | -2.71 | -2.40 |
|  | Role[Parent] | 1.57 | 0.11 | 1.35 | 1.78 |
| | Random effect: $\sigma_1\vert _{\text{Family ID}}$ | 0.55 | 0.09 | 0.39 | 0.72 |
| Phase (rad) | Intercept | 1.82 | 2.05 | -2.43 | 5.46 |
|  | Role[Parent] | -3.39 | 1.73 | -7.70 | -0.70 |
| | Random effect: $\sigma_1\vert _{\text{Family ID}}$ | 11.36 | 1.75 | 7.93 | 14.71 |

**Table S5.** Statistical analysis of LIDS parameters (cycle length and phase) in infants at 3, 6 and 12 months. Bayesian generalized mixed model analysis with random intercept per infant and time point: estimated mean, standard deviation and limits of the credible interval of posterior distributions for fixed and random effects. Statistically significant effects are shaded in gray. Reference levels for categorical fixed effects: 3 months for time point, female for sex and separate room for sleep location. Abbreviations: Std, standard deviation; HDI, highest density interval.

| LIDS parameter |  | Mean | Std. | 95% HDI:<br>lower limit | 95% HDI:<br>upper limit |
| --- | --- | --- | --- | --- | --- |
| Cycle length (min) | Intercept | 52.14 | 4.89 | 42.82 | 62.04 |
|  | Time point[6 months] | -0.08 | 0.69 | -1.45 | 1.26 |
|  | Time point[12 months] | 4.70 | 0.93 | 2.85 | 6.51 |
|  | Sex[Male] | 1.34 | 0.74 | -0.08 | 2.83 |
|  | Bout duration (h) | 1.01 | 0.07 | 0.87 | 1.15 |
|  | Night sleep (% , x10) | 0.18 | 0.66 | -1.13 | 1.47 |
|  | Sleep location[Separate bed] | 0.97 | 0.72 | -0.43 | 2.40 |
|  | Sleep location[Attached cot] | 0.61 | 1.06 | -1.49 | 2.66 |
|  | Sleep location[Parents' bed] | 2.94 | 0.89 | 1.16 | 4.64 |
| | $\sigma_1$\|Subject ID | 3.64 | 0.36 | 2.94 | 4.34 |
| | $\sigma_{\text{Time point}}$\|Subject ID[6 months] | 2.72 | 0.99 | 0.60 | 4.62 |
| | $\sigma_{\text{Time point}}$\|Subject ID[12 months] | 3.11 | 0.82 | 1.46 | 4.73 |
| Phase (rad) | Intercept | 10.32 | 19.72 | -32.63 | 46.16 |
|  | Time point[6 months] | -2.54 | 2.81 | -8.05 | 2.39 |
|  | Time point[12 months] | -4.18 | 3.64 | -10.93 | 3.59 |
|  | Sex[Male] | 1.27 | 2.36 | -3.41 | 5.92 |
|  | Bout duration (h) | -1.13 | 0.24 | -1.63 | -0.65 |
|  | Night sleep (% , x10) | 0.45 | 2.39 | -3.84 | 5.79 |
|  | Sleep location[Separate bed] | 1.16 | 2.67 | -3.76 | 6.54 |
|  | Sleep location[Attached cot] | -1.04 | 4.50 | -9.64 | 8.13 |
|  | Sleep location[Parents' bed] | 0.66 | 3.02 | -5.49 | 6.95 |
| | $\sigma_1$\|Subject ID | 7.34 | 4.46 | 0.00 | 16.27 |
| | $\sigma_{\text{Time point}}$\|Subject ID[6 months] | 3.02 | 1.88 | 0.00 | 6.27 |
| | $\sigma_{\text{Time point}}$\|Subject ID[12 months] | 2.78 | 2.22 | 0.00 | 7.33 |

**Table S6.** Statistical analysis of LIDS parameters (amplitude, offset and slope) in infants at 3, 6 and 12 months. Bayesian generalized mixed model analysis with random intercept per infant and time point: estimated mean, standard deviation and limits of the credible interval of posterior distributions for fixed and random effects. Statistically significant effects are shaded in gray. Reference levels for categorical fixed effects: 3 months for time point, female for sex and separate room for sleep location. Abbreviations: Std, standard deviation; HDI, highest density interval.

| LIDS parameter |  | Mean | Std. | 95% HDI:<br>lower limit | 95% HDI:<br>upper limit |
| --- | --- | --- | --- | --- | --- |
| Amplitude (LIDS unit) | Intercept | 18.98 | 1.69 | 15.63 | 22.27 |
|  | Time point[6 months] | -2.27 | 0.23 | -2.71 | -1.83 |
|  | Time point[12 months] | -1.57 | 0.32 | -2.19 | -0.93 |
|  | Sex[Male] | -0.28 | 0.33 | -0.92 | 0.38 |
|  | Bout duration (h) | -0.61 | 0.02 | -0.64 | -0.57 |
|  | Night sleep (% , x10) | 0.03 | 0.23 | -0.41 | 0.49 |
|  | Sleep location[Separate bed] | -0.21 | 0.27 | -0.73 | 0.31 |
|  | Sleep location[Attached cot] | -0.72 | 0.37 | -1.45 | -0.02 |
|  | Sleep location[Parents' bed] | -0.90 | 0.33 | -1.53 | -0.27 |
| | $\sigma_1$\|Subject ID | 1.88 | 0.13 | 1.63 | 2.15 |
| | $\sigma_{\text{Time point}}$\|Subject ID[6 months] | 1.43 | 0.16 | 1.12 | 1.73 |
| | $\sigma_{\text{Time point}}$\|Subject ID[12 months] | 1.57 | 0.17 | 1.24 | 1.91 |
| Offset (LIDS unit) | Intercept | 77.03 | 3.27 | 70.81 | 83.58 |
|  | Time point[6 months] | 3.22 | 0.44 | 2.38 | 4.10 |
|  | Time point[12 months] | 1.50 | 0.63 | 0.27 | 2.72 |
|  | Sex[Male] | -0.79 | 0.51 | -1.77 | 0.21 |
|  | Bout duration (h) | 0.93 | 0.05 | 0.83 | 1.02 |
|  | Night sleep (% , x10) | -0.07 | 0.45 | -0.94 | 0.81 |
|  | Sleep location[Separate bed] | 0.14 | 0.50 | -0.83 | 1.14 |
|  | Sleep location[Attached cot] | 0.14 | 0.71 | -1.20 | 1.56 |
|  | Sleep location[Parents' bed] | 0.81 | 0.59 | -0.32 | 2.02 |
| | $\sigma_1$\|Subject ID | 2.55 | 0.23 | 2.13 | 3.00 |
| | $\sigma_{\text{Time point}}$\|Subject ID[6 months] | 1.30 | 0.55 | 0.08 | 2.21 |
| | $\sigma_{\text{Time point}}$\|Subject ID[12 months] | 1.73 | 0.57 | 0.50 | 2.81 |
| Slope (LIDS unit/h) | Intercept | -3.32 | 0.68 | -4.61 | -1.99 |
|  | Time point[6 months] | 0.65 | 0.09 | 0.46 | 0.83 |
|  | Time point[12 months] | 0.83 | 0.12 | 0.60 | 1.06 |
|  | Sex[Male] | -0.06 | 0.09 | -0.23 | 0.12 |
|  | Bout duration (h) | 0.15 | 0.01 | 0.12 | 0.17 |
|  | Night sleep (% , x10) | -0.03 | 0.09 | -0.21 | 0.15 |
|  | Sleep location[Separate bed] | 0.11 | 0.09 | -0.08 | 0.29 |
|  | Sleep location[Attached cot] | 0.17 | 0.14 | -0.11 | 0.46 |
|  | Sleep location[Parents' bed] | 0.34 | 0.11 | 0.13 | 0.56 |
| | $\sigma_1$\|Subject ID | 0.37 | 0.05 | 0.28 | 0.46 |
| | $\sigma_{\text{Time point}}$\|Subject ID[6 months] | 0.29 | 0.12 | 0.03 | 0.48 |
| | $\sigma_{\text{Time point}}$\|Subject ID[12 months] | 0.12 | 0.08 | 0.00 | 0.28 |

**Table S7.** Relationship between infant LIDS cycle length and bout duration - statistical analysis without accounting for sleep bout duration. Bayesian generalized mixed model analysis of infant LIDS cycle length with random intercept per infant: estimated mean, standard deviation and limits of the credible interval of posterior distributions for fixed and random effects. Statistically significant effects are shaded in gray. Reference levels for categorical fixed effects: 3 months for time point, female for sex and separate room for sleep location. Abbreviations: Std, standard deviation and HDI, highest density interval.

| LIDS parameter |  | Mean | Std. | 95% HDI:<br>lower limit | 95% HDI:<br>upper limit |
| --- | --- | --- | --- | --- | --- |
| Cycle length (min) | Intercept | 53.68 | 5.02 | 43.81 | 63.48 |
|  | Time point[6 months] | 1.21 | 0.73 | -0.22 | 2.64 |
|  | Time point[12 months] | 7.62 | 0.94 | 5.78 | 9.46 |
|  | Sex[Male] | 0.70 | 0.76 | -0.76 | 2.21 |
|  | Night sleep (% , x10) | 0.71 | 0.68 | -0.65 | 2.04 |
|  | Sleep location[Separate bed] | 0.50 | 0.74 | -0.94 | 1.97 |
|  | Sleep location[Attached cot] | 0.30 | 1.08 | -1.86 | 2.39 |
|  | Sleep location[Parents' bed] | 2.07 | 0.90 | 0.31 | 3.85 |
| | $\sigma_1$\|Subject ID | 3.67 | 0.38 | 2.96 | 4.44 |
| | $\sigma_{\text{Time point\vert Subject ID [6 months]}}$ | 3.38 | 0.97 | 1.47 | 5.36 |
| | $\sigma_{\text{Time point\vert Subject ID [12 months]}}$ | 3.24 | 0.86 | 1.51 | 4.93 |

**Table S8.** Relationship between infant LIDS cycle length and sleep bout duration - statistical analysis of the first 3 hours of long bouts. Bayesian generalized mixed model analysis of infant LIDS cycle length in bouts *>*5 hours based on parameter estimates for the first 3h of each sleep bout (*n* = 4357 bouts): estimated mean, standard deviation and limits of the credible interval of posterior distributions for fixed and random effects. Statistically significant effects are shaded in gray. Reference levels for categorical fixed effects: 3 months for time point, female for sex and separate room for sleep location. Abbreviations: Std, standard deviation; HDI, highest density interval.

| LIDS parameter |  | Mean | Std. | 95% HDI:<br>lower limit | 95% HDI:<br>upper limit |
| --- | --- | --- | --- | --- | --- |
| Cycle length (min) | Intercept | 57.04 | 4.59 | 48.03 | 66.01 |
|  | Time point[6 months] | -0.96 | 0.61 | -2.16 | 0.21 |
|  | Time point[12 months] | 3.05 | 0.83 | 1.46 | 4.68 |
|  | Sex[Male] | -0.09 | 0.68 | -1.41 | 1.25 |
|  | Bout duration (h) | -0.16 | 0.08 | -0.31 | -0.01 |
|  | Night sleep (% , x10) | 0.56 | 0.62 | -0.67 | 1.77 |
|  | Sleep location[Separate bed] | 0.18 | 0.65 | -1.10 | 1.43 |
|  | Sleep location[Attached cot] | 0.19 | 0.97 | -1.79 | 2.03 |
|  | Sleep location[Parents' bed] | 0.86 | 0.79 | -0.65 | 2.45 |
| | $\sigma_1$\|Subject ID | 3.31 | 0.31 | 2.71 | 3.93 |
| | $\sigma_{\text{Time point}}$\|Subject ID [6 months] | 1.02 | 0.67 | 0.00 | 2.21 |
| | $\sigma_{\text{Time point}}$\|Subject ID [12 months] | 1.01 | 0.69 | 0.00 | 2.27 |

**Table S9.** Statistical analysis of infant LIDS parameters at 12 months including breastfeeding status. Bayesian generalized mixed model analysis with random intercept per infant: estimated mean, standard deviation and limits of the credible interval of posterior distributions for fixed and random effects. Statistically significant effects are shaded in gray. Reference levels for categorical fixed effects: female for sex, no for breastfed and separate room for sleep location. Abbreviations: Std, standard deviation; HDI, highest density interval.

| LIDS parameter |  | Mean | Std. | 95% HDI:<br>lower limit | 95% HDI:<br>upper limit |
| --- | --- | --- | --- | --- | --- |
| Cycle length (min) | Intercept | 41.22 | 10.06 | 21.13 | 60.88 |
|  | Sex[Male] | 0.58 | 0.96 | -1.25 | 2.51 |
|  | Bout duration (h) | 1.00 | 0.11 | 0.78 | 1.22 |
|  | Breastfed[Yes] | 2.47 | 1.04 | 0.41 | 4.49 |
|  | Night sleep (% , x10) | 1.92 | 1.21 | -0.46 | 4.33 |
|  | Sleep location[Separate bed] | 1.13 | 1.17 | -1.09 | 3.50 |
|  | Sleep location[Parents' bed] | 3.15 | 1.33 | 0.57 | 5.80 |
| | $\sigma_1$\|Subject ID | 3.89 | 0.59 | 2.80 | 5.09 |
| Amplitude (LIDS unit) | Intercept | 16.61 | 3.61 | 9.77 | 23.93 |
|  | Sex[Male] | -0.28 | 0.34 | -0.95 | 0.37 |
|  | Bout duration (h) | -0.38 | 0.03 | -0.43 | -0.32 |
|  | Breastfed[Yes] | 0.02 | 0.37 | -0.71 | 0.73 |
|  | Night sleep (% , x10) | -0.13 | 0.44 | -0.98 | 0.73 |
|  | Sleep location[Separate bed] | 0.27 | 0.41 | -0.55 | 1.07 |
|  | Sleep location[Parents' bed] | -1.11 | 0.46 | -2.03 | -0.22 |
| | $\sigma_1$\|Subject ID | 1.71 | 0.14 | 1.44 | 1.99 |
| Offset (LIDS unit) | Intercept | 80.72 | 6.05 | 69.18 | 92.85 |
|  | Sex[Male] | -0.67 | 0.56 | -1.79 | 0.42 |
|  | Bout duration (h) | 0.31 | 0.06 | 0.19 | 0.43 |
|  | Breastfed[Yes] | 1.40 | 0.62 | 0.18 | 2.60 |
|  | Night sleep (% , x10) | 0.40 | 0.73 | -1.08 | 1.78 |
|  | Sleep location[Separate bed] | -0.88 | 0.70 | -2.32 | 0.43 |
|  | Sleep location[Parents' bed] | -0.68 | 0.79 | -2.24 | 0.83 |
| | $\sigma_1$\|Subject ID | 2.60 | 0.26 | 2.10 | 3.10 |
| Slope (LIDS unit/h) | Intercept | -1.63 | 1.18 | -3.92 | 0.69 |
|  | Sex[Male] | -0.16 | 0.11 | -0.37 | 0.06 |
|  | Bout duration (h) | 0.12 | 0.02 | 0.08 | 0.16 |
|  | Breastfed[Yes] | -0.12 | 0.12 | -0.35 | 0.11 |
|  | Night sleep (% , x10) | -0.10 | 0.14 | -0.38 | 0.19 |
|  | Sleep location[Separate bed] | 0.10 | 0.13 | -0.17 | 0.35 |
|  | Sleep location[Parents' bed] | 0.51 | 0.15 | 0.20 | 0.81 |
| | $\sigma_1$\|Subject ID | 0.44 | 0.06 | 0.33 | 0.56 |
| Phase (rad) | Intercept | 7.82 | 21.38 | -32.37 | 49.83 |
|  | Sex[Male] | 0.58 | 1.82 | -3.00 | 3.82 |
|  | Bout duration (h) | -1.41 | 0.52 | -2.27 | -0.84 |
|  | Breastfed[Yes] | 1.18 | 2.84 | -4.52 | 6.60 |
|  | Night sleep (% , x10) | 0.51 | 2.43 | -4.04 | 5.48 |
|  | Sleep location[Separate bed] | -0.41 | 2.22 | -4.61 | 4.09 |
|  | Sleep location[Parents' bed] | 0.88 | 3.30 | -5.39 | 7.73 |
| | $\sigma_1$\|Subject ID | 4.46 | 6.52 | 1.78 | 5.88 |

**Table S10.**
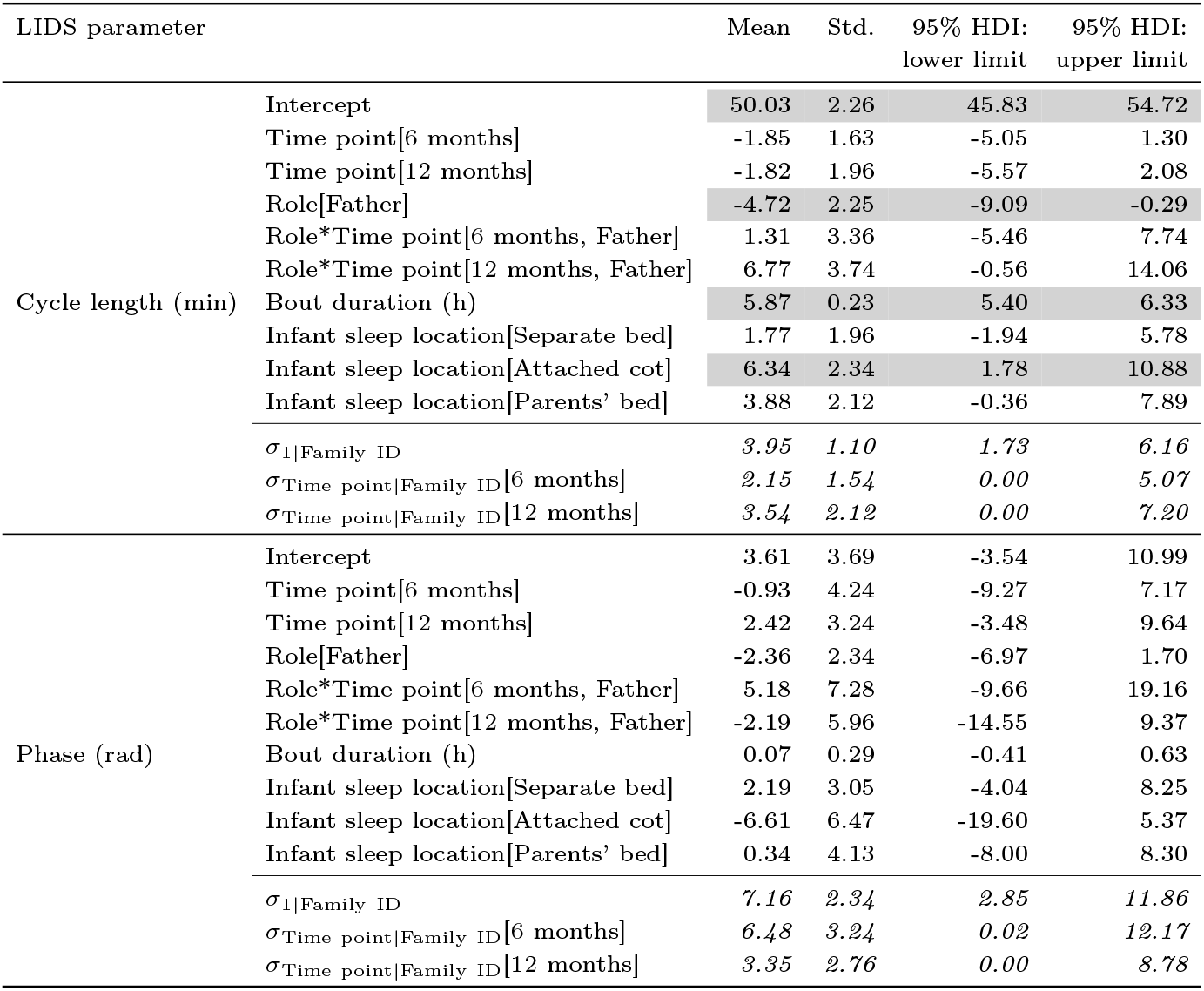
Statistical analysis of parental LIDS parameters (cycle length and phase) across time points. Bayesian generalized mixed model analysis with random intercept by family and random slopes per time point: estimated mean, standard deviation and limits of the credible interval of posterior distributions for fixed and random effects. Statistically significant effects are shaded in gray. Reference levels for categorical fixed effects: 3 months for time point, mother for role and separate room for sleep location. Abbreviations: Std, standard deviation; HDI, highest density interval.

**Table S11.** Statistical analysis of parental LIDS parameters (amplitude, offset and slope) across time points. Bayesian generalized mixed model analysis with random intercept by family and random slopes per time point: estimated mean, standard deviation and limits of the credible interval of posterior distributions for fixed and random effects. Statistically significant effects are shaded in gray. Reference levels for categorical fixed effects: 3 months for time point, mother for role and separate room for sleep location. Abbreviations: Std, standard deviation; HDI, highest density interval.

| LIDS parameter |  | Mean | Std. | 95% HDI:<br>lower limit | 95% HDI:<br>upper limit |
| --- | --- | --- | --- | --- | --- |
| Amplitude (LIDS unit) | Intercept | 9.44 | 0.33 | 8.79 | 10.08 |
|  | Time point[6 months] | -0.24 | 0.21 | -0.66 | 0.16 |
|  | Time point[12 months] | -0.28 | 0.26 | -0.80 | 0.23 |
|  | Role[Father] | -0.50 | 0.23 | -0.94 | -0.07 |
|  | Role*Time point[6 months, Father] | 0.51 | 0.33 | -0.15 | 1.14 |
|  | Role*Time point[12 months, Father] | 0.63 | 0.37 | -0.10 | 1.35 |
|  | Bout duration (h) | -0.27 | 0.02 | -0.31 | -0.22 |
|  | Infant sleep location[Separate bed] | 0.26 | 0.30 | -0.34 | 0.83 |
|  | Infant sleep location[Attached cot] | 0.65 | 0.37 | -0.06 | 1.41 |
|  | Infant sleep location[Parents' bed] | 1.38 | 0.35 | 0.68 | 2.05 |
| | $\sigma_{1\vert Family\ ID}$ | 1.25 | 0.14 | 0.99 | 1.53 |
| | $\sigma_{Time\ point\vert Family\ ID}$ [6 months] | 1.08 | 0.20 | 0.70 | 1.47 |
| | $\sigma_{Time\ point\vert Family\ ID}$ [12 months] | 1.09 | 0.20 | 0.72 | 1.50 |
| Offset (LIDS unit) | Intercept | 90.30 | 0.63 | 89.04 | 91.51 |
|  | Time point[6 months] | -0.27 | 0.35 | -0.94 | 0.44 |
|  | Time point[12 months] | -1.07 | 0.46 | -1.98 | -0.16 |
|  | Role[Father] | -1.17 | 0.50 | -2.13 | -0.19 |
|  | Role*Time point[6 months, Father] | -1.45 | 0.74 | -2.89 | -0.02 |
|  | Role*Time point[12 months, Father] | -0.30 | 0.83 | -1.93 | 1.34 |
|  | Bout duration (h) | 0.22 | 0.06 | 0.11 | 0.33 |
|  | Infant sleep location[Separate bed] | -0.28 | 0.52 | -1.29 | 0.71 |
|  | Infant sleep location[Attached cot] | -0.09 | 0.62 | -1.30 | 1.14 |
|  | Infant sleep location[Parents' bed] | -1.63 | 0.59 | -2.81 | -0.51 |
| | $\sigma_{1\vert Family\ ID}$ | 2.19 | 0.26 | 1.71 | 2.71 |
| | $\sigma_{Time\ point\vert Family\ ID}$ [6 months] | 0.59 | 0.38 | 0.00 | 1.27 |
| | $\sigma_{Time\ point\vert Family\ ID}$ [12 months] | 0.98 | 0.51 | 0.00 | 1.82 |
| Slope (LIDS unit/h) | Intercept | -0.97 | 0.18 | -1.33 | -0.61 |
|  | Time point[6 months] | 0.19 | 0.10 | -0.02 | 0.39 |
|  | Time point[12 months] | 0.19 | 0.12 | -0.05 | 0.44 |
|  | Role[Father] | 0.09 | 0.14 | -0.18 | 0.36 |
|  | Role*Time point[6 months, Father] | -0.05 | 0.21 | -0.47 | 0.35 |
|  | Role*Time point[12 months, Father] | -0.12 | 0.23 | -0.55 | 0.36 |
|  | Bout duration (h) | 0.04 | 0.02 | 0.01 | 0.08 |
|  | Infant sleep location[Separate bed] | -0.10 | 0.13 | -0.35 | 0.16 |
|  | Infant sleep location[Attached cot] | -0.21 | 0.16 | -0.53 | 0.10 |
|  | Infant sleep location[Parents' bed] | -0.01 | 0.15 | -0.30 | 0.29 |
| | $\sigma_{1\vert Family\ ID}$ | 0.44 | 0.06 | 0.32 | 0.57 |
| | $\sigma_{Time\ point\vert Family\ ID}$ [6 months] | 0.15 | 0.10 | 0.00 | 0.33 |
| | $\sigma_{Time\ point\vert Family\ ID}$ [12 months] | 0.21 | 0.13 | 0.00 | 0.43 |

**Table S12.**
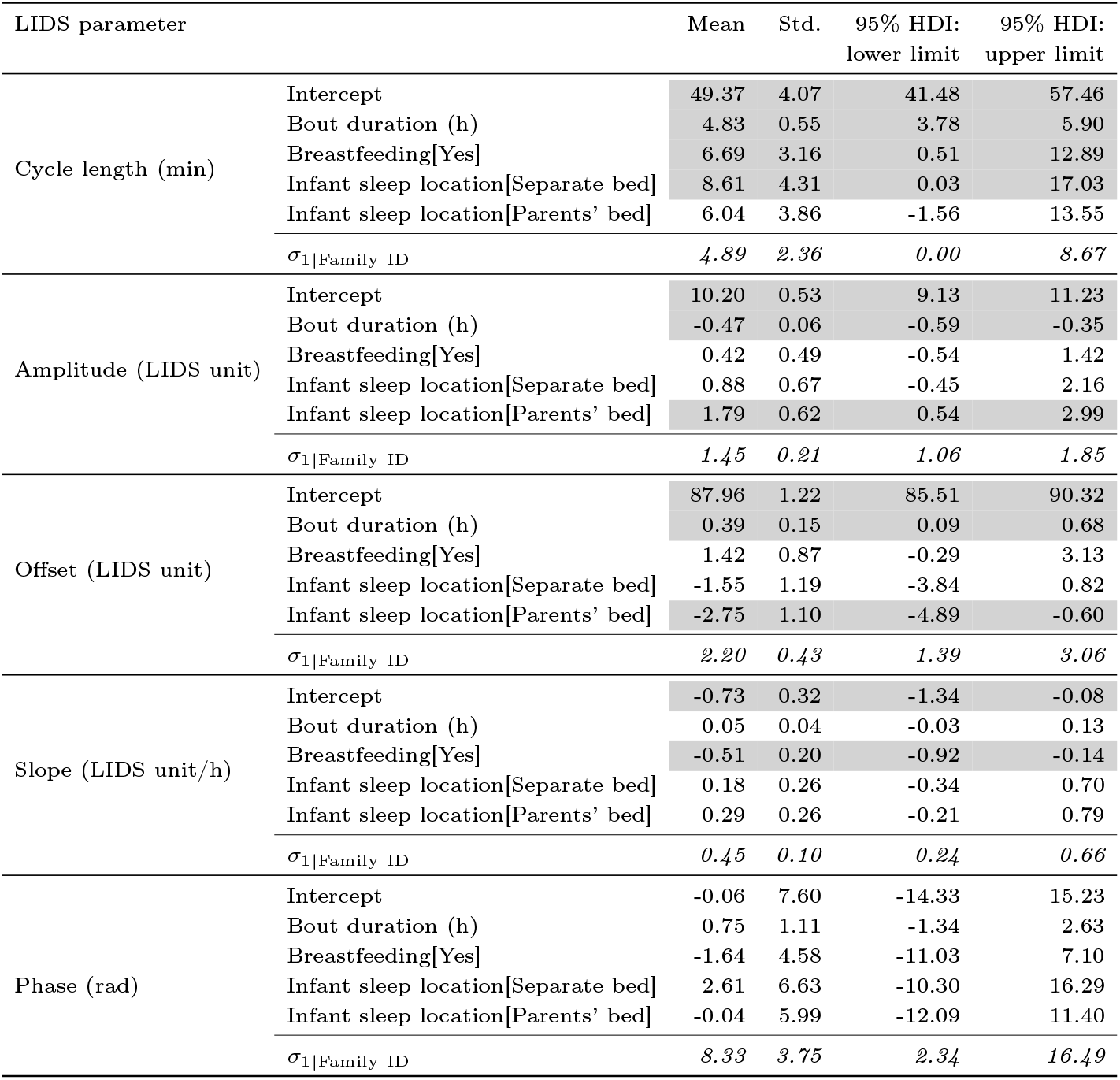
Statistical analysis of mothers’ LIDS parameters at 12 months including breastfeeding status. Bayesian generalized mixed model analysis with random intercept per family: estimated mean, standard deviation and limits of the credible interval of posterior distributions for fixed and random effects. Statistically significant effects are shaded in gray. Reference levels for categorical fixed effects: no for breastfeeding and separate room for sleep location. Abbreviations: Std, standard deviation; HDI, highest density interval.

